# Development and characterization of bioinks for 3D bioprinting of in vitro skeletal muscle constructs

**DOI:** 10.1101/2024.11.01.621422

**Authors:** Rodi Kado Abdalkader, Kosei Yamauchi, Satoshi Konishi, Takuya Fujita

**Affiliations:** Ritsumeikan Global-Innovation Research Organization (R-GIRO), Ristumeikan University, 1-1-1 Noji-Higashi, Kusatsu, Shiga, 525-8577 Japan; Department of Mechanical Engineering, Ritumeikan University, 1-1-1 Noji-Higashi, Kusatsu, Shiga, 525-8577 Japan; Department of Pharmaceutical Sciences, Ritsumeikan University, 1-1-1 Noji-Higashi, Kusatsu, Shiga, 525-8577 Japan

## Abstract

The use of 3D bioprinting to construct *in vitro* skeletal muscle models presents a promising approach; however, selecting an optimal bioink remains a common challenge. This study focuses on the development and characterization of bioinks for extrusion-based 3D bioprinting, specifically targeting the creation of accurate skeletal muscle models. By exploring various compositions of alginate, gelatin, fibrinogen, and nanofiber cellulose, we evaluate these formulations based on printability and their support for the growth and differentiation of C2C12 myoblast cells.

While alginate provided a strong, stable matrix for printing scaffolds embedded with C2C12 cells, it did not effectively promote cell growth and differentiation. The addition of fibrinogen to alginate enhanced cell growth and differentiation but was limited mainly to the scaffold surfaces, even with the inclusion of gelatin as a sacrificial ink. Notably, replacing alginate with nanofiber cellulose (NFC) alongside fibrinogen significantly improved cell growth and differentiation, leading to the formation of mature myotubes. Cell distribution was observed both inside and on the surfaces of the scaffolds, indicating effective spatial cell distribution. Furthermore, the scaffolds were tailored to form skeletal muscle bundles anchored between PDMS pillars for contractility testing. Upon exposure to electrical stimulation, the cells displayed measurable displacement, demonstrating contractile function.

These findings offer valuable insights into optimizing bioink formulations that promote myoblast growth and differentiation into skeletal muscle *in vitro*, with potential applications in future neuromuscular disease modeling.

## 1. Introduction

Muscular diseases, including muscle atrophy, represent significant challenges for medical treatment and management, highlighting a critical need for the development of advanced preclinical models that can accurately mimic neuromuscular diseases^1^. This is essential for enhancing drug discovery processes and developing new therapeutic strategies. Traditionally, animal models have been the cornerstone of disease modeling and drug discovery. They can offer valuable insights into disease mechanisms, progression, and therapeutic responses. However, notable physiological, genetic, and immunological differences between animals and humans often result in variation in disease manifestation and drug efficacy, which may not translate effectively to human conditions^2^.

To bridge these gaps, recent research advances have focused on developing new *in vitro* models that utilize human cells and tissues, such as organ-on-a-chip (OoC) technology^3^. These models offer a more relevant three-dimensional representation of human tissues and can emulate the biomechanical environment of organs, thereby providing a more accurate platform for disease modeling and drug testing^4, 5^.

A particularly promising avenue in this field is 3D bioprinting, which facilitates the fabrication of complex, layered structures of a tissue using techniques such extrusion based bioprinting^6^. This process utilizes bioinks—biocompatible gels composed of natural or synthetic extracellular matrix (ECM) materials mixed with cells^7^. Among the bioinks employed, alginate is favored for its ease of gelation through ionic cross-linking with calcium chloride, though it lacks the adhesion molecules necessary for inducing interaction with cells^8^. Conversely, collagen and its derivative gelatin, which are biologically relevant to the human ECM, pose technical challenges during printing due to their thermosensitivity that require precise conditions for effective printing^9^. Moreover, the mechanical properties and cellular response are highly dependent on the concentration, type of collagen used, and cross-linking method which can affect the fate of the printed tissue^10^. An alternative strategy involves the use of fibrin gel, known for its biological competency and ability to support the differentiation and growth of various cell types, such as endothelial and muscles cells^11,12^. However, fibrinogen’s poor printability prior cross-linking limits its use in bioprinting^13^. Combining fibrinogen with polysaccharides or nanocellulose thickener has shown potential in overcoming these limitations, providing a composite bioink that supports robust cell growth while maintaining good printability and structural integrity^14,15^. Yet, it remains uncertain whether these bioink formulations can sufficiently support the growth and differentiation of skeletal muscle cells under 3D culturing conditions. Further investigation is required to optimize these bioinks for muscle tissue engineering, ensuring they can mimic the complex environment necessary for muscle cell maturation and functionality.

In this study, we aimed to assess the potential of using a combination of fibrinogen gel with polysaccharides or nanofiber cellulose bioinks. We examined the printability, and compatibility of these bioinks with murine myoblast cells C2C12 using an extrusion-based bioprinter and then evaluated their differentiation and functions. Our approach provides valuable insights into the development of advanced 3D models of skeletal muscles for potential application in neuro-musculoskeletal diseases modeling.

## 2. Materials and methods

### 2.1. C2C12 cell culturing

C2C12 mouse myoblast cells, obtained from RIKEN Bioresource Research Centre in Japan, were cultured in DMEM supplemented with 10% fetal bovine serum. Cell passaging was carried out using a trypsin-EDTA solution at a subculture ratio of 1:10. To induce differentiation into myotubes, the culturing medium was changed to DMEM high glucose (Wako, Osaka, Japan) supplemented with 5% horse serum and 100 ng/ mL insuline growth factor (IGF) (Wako, Osaka, Japan).

### 2.2. Bioink preparation

The formulation of alignate-gelatin bioinks involved a stepwise procedure. Initially, sodium alginate (Wako, Osaka, Japan), gelatin from procine (Nitta Gelatin, Osaka, Japan), and glycerol was added into PBS while being subjected to magnetic stirring at 60 °C for a duration of 30 minutes. The PH was adjusted with 0.1 M NaOH and confirmed with a paper-based PH kit until the PH value was neutralized. Prior to use, the bioink was warmed at 37 °C and then centrifued at 1500 rmp for 5 min to remove bubbles. For the formulation of alignate-fibrinogen and nano fiber cellulose (NFC) (3D-NanoFibGrow-I (3D NF); synthesized by acetic acid bacteria using sugar beet) (Nano T-Sailing, LLC. Tokushima, Japan) was used in the bioinks. Sodium alginate or NFC was gradually introduced into PBS while being subjected to magnetic stirring at 50 °C. That followed by the addtion of gelatine, hyaluronic acid (Wako, Osaka, Japan), and glycerin under magnetic stirring at 50 °C for 60 minutes. Finally, fibrinogen derived from bovine (Wako, Osaka, Japan) was added to the mixure at 37 °C and mixed well. The bioink solutions were carefully transferred into 10 mL syringes and stored for a maximum period of 1week at 4 °C until their intended use (**Table. 1**).

### 2.3. Design of scaffolds and 3D bioprinting

To print scaffolds, a custom-made round shape or square shape with dimensions of 9mm x 9mm x0.8 mm (LxWxH) and 14mm x14mm x 0.8mm (LxWxH), respectively were designed using Tinkercad (https://www.tinkercad.com). The design was then converted into STL files and sliced and processed using heartware software. The pritability of bioinks was determined using a 3D bioprinter (INKRIDIBLE, Cellink, Sweden) with cartridges loaded with bioinks. Printing was performed at a temperature of 25°C. The pritability of each bioink was evaluated by varying the printing speeds (4 mm/s, 8 mm/2, 10 mm/s) and pneumatic air pressure (10 kPa, 20 kPa, 30 kPa, and 40 kPa) using a 22 G printing nozzle (Cellink, Sweden). To prepare the bioink mixture for bioprinting C2C12 cells, we mixed 1×10^7^ cells in 0.3 mL of cell culturing medium with 3 mL of the bioink using a cellmixer (Cellink, Sweden) or cells were directly mixed with the bioink and then trnasferred to the 3 mL printing cartiridge (Cellink, Sweden). After printing, the scaffolds were cross-linked with 0.5 M CaCl2 for 3 minutes and washed with Hanks buffer containing calicum (HBSS+). Scaffolds were then immersed in cell culturing medium supplemented with 5 U/mL thrombin (Wako, Osaka, Japan) and placed at 37°C with 5% CO^2^ for 60 minutes and then replaced by DMEM medium supplemented with 10% fetal bovine serum and 1μg/mL aprotinin (Wako, Osaka, Japan) for 3 days and then replaced by the differentiation medium comprised of DMEM high glucose supplemented with 5% horse serum and 1μg/mL aprotinin. The culturing media were change on a daily-base. The fidelity of the printing was assessed by measuring the printing area using imageJ software acorrding the equation.

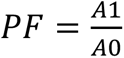

In this equation, *PF*, *A1*, and *A0* represent printing fidelity, the sample’s actual area, and the sample’s theoritical area.

### 2.4. Design and fabrication of pillar substrates for 3D bioprinting

To fabricate polydimethylsiloxane (PDMS) pillars and substrates, Sylgard 184 PDMS two-part elastomer (Dow Corning Corporation, Midland, MI, USA) was mixed in a 10:1 ratio of pre-polymer to curing agent. The mixture was poured into molds with heights of 3 mm and 8 mm, respectively. The PDMS molds were then degassed and cured at 65 °C for 6 hours. After curing, the PDMS was removed from the molds, trimmed, and cut into 10×10 mm sheets using a blade. Pillars were created by using a 1 mm punch on the 8 mm-thick PDMS blocks. Each pillar was then affixed to the PDMS sheet, with a 3 mm gap between each pillar, using the PDMS curing agent. The assembly was cured again at 65 °C for 6 hours. Before use, the PDMS pillar substrates were immersed in 2% Pluronic F127 (Biotium, USA) overnight at 4 °C and then sterilized by UV exposure for 30 minutes at room temperature. To print scaffolds, two custom-made; a rectangular structure and interlocking rings with dimensions of 8mmx 4mmx0.8 mm (LxWxH) and 7.4 mmx 4mm x 0.8mm (LxWxH), respectively, were designed using Tinkercad and processed similarly using heartware softare. G-codes can be found in **supplemental data 1**.

### 2.5. Cell viability

Live-dead cell analysis was conducted using the LIVE/DEAD Cell Imaging Kit (Thermo Fisher). For Calcein AM and Hoechst staining, cells were incubated with Calcein AM and Hoechst 33342 (Dojindo) at a final concentration of 10 µg/mL in HBSS+ at 37°C with 5% CO2 for 30 minutes. Subsequently, images of the cells were captured using a fluorescence microscope (KEYENCE, Tokyo, Japan).

### 2.6. Electrical stimulus and microscopy imaging

For the electrical stimulus test, carbon electrodes in a diameter of Φ1.0 mm were employed. The probes were fixed in the cell culturing dish leaving a space of 1 cm and then connected to an electricity generator (FGX-295, Texio) while controlled by a PC using LabVIEW 2017 software (National Instruments Corporation). Parameters were adjusted to a pulse of 2 ms, intensity of 5 V, and frequency of 1Hz. The electrical pulse stimulus (EPS) was adjusted to one stimulus per second. The contraction of the cells constructs was observed by inverted microscopy (Olympus) and then analysed with imageJ software.

### 2.7. 3D microscopic imaging and structure analysis

Scaffolds were washed with PBS twice and then fixed with 4% paraformaldehyde in PBS-for 60 minutes at 25°C or in methanol at −20 °C for 15 minutes with alginate scaffolds. Samples were then permeabilized with 0.1% Triton X-100 at 4 °C for overnight. Subsequently, scaffolds were incubated with ActinGreen 488 or ActinGreen 555 (Thermo Fisher Scientific) at 4 °C for overnight. And then scaffolds were washed with PBS twice and then incubated with Hoechst 33342 (Dojindo Molecular Technologies, Inc.) in a final concentration of 10 µg/ mL for 60 minutes at 25°C. For imaging, we used confocal scanning microscope (FV3000; Olympus) and z-scans of 2-3 µm were obtained from each scaffold. Images were then analysed using FV31S-SW software. For the investigation of myotubes orientation, the Directionality function analysis as well as OrientationJ in ImageJ software was employed (National Institute of Health, Maryland, USA).

### 2.8. Data visualization and statistics

The unpaired t-test and Tukey’s test were performed using GraphPad prism 8 (GraphPad Software, La Jolla California, USA). Data mining and visualization was conducted by Orange 3 software (Version 3.23.1; Bioinformatics Laboratory, Faculty of Computer and Information Science, University of Ljubljana, Slovenia, Python Jupyter notebook 6.1.4, with Pandas and Bioinfokit packages, and matplotlib and seaborn packages.

## 3. Results

### 3.1. Impact of gelatin concentration on printing fidelity and cell encapsulation in alginate-based bioinks

Gelatin is well known and bioactive component as well as a sacrificial agent that can be released upon incubation at 37 °C allowing for nutrient to be diffused deeper into scaffolds and thus enhance their survive. To evaluate how gelatine composition affects the print quality, we tested varying amounts of gelatin with a fixed portion of sodium alginate (**Table 1**). As shown in **Figure 1A**, the printed grid-like structures varied significantly across bioinks. Bioink BI_1, which lacked gelatine, produced less defined structures with irregular line thickness. As the gelatine concentration increased, the print fidelity improved, with bioink BI_2 (3% w/v gelatine) yielding the most uniform lines. The differences in print fidelity were further confirmed through microscopic analysis (**Figure 1B**). Bioinks BI_1 and BI_4, which lacked or had low concentrations of gelatine, produced lines with irregular widths and poorly defined intersections. Bioinks BI_2 and BI_3 with higher gelatin concentrations, showed improved line quality and better-defined intersections. Quantitative analysis (**Figure 1C**) showed that line width significantly decreased as gelatine concentration increased (p < 0.0001) between bioinks BI_1 and BI_2 compared to BI_3 and BI_4). However, there was no significant difference in the intersection angles between Bioinks BI_1 and BI_4. To test cell survival in alginate-gelatin bioinks, we selected bioink BI_3 and encapsulated C2C12 cells for evaluation of their survival. While the cells could survive and be remined viable within the constructs for 24 hours (**Figure 1D; (I)**), there was no evidence of cell elongation during 12 day of cell culturing, which typically indicates differentiation into myotubes (**Figure 1D; (II)**).

**Table 1.** The formulation of bioinks.

| **Bioink** | **Sodium Alginate**<br>**% (w/v)** | **Gelatine**<br>**% (w/v)** | **Hyaluronic acid**<br>**% (w/v)** | **Fibrinogen**<br>**(mg/mL)** | **Glycerin**<br>**% (v/v)** | **NFC**<br>**% (v/v)** |
| --- | --- | --- | --- | --- | --- | --- |
| **BI_1** | 3.5 | 0 | — | — |  |  |
| **BI_2** | 3.5 | 3 | — | — |  |  |
| **BI_3** | 3.5 | 1.5 | — | — |  |  |
| **BI_4** | 3.5 | 0.75 | — |  |  |  |
| **BI_5** | 3.5 | 0.5 | — | 10 | 0.5 |  |
| **BI_6** | 3.5 | 0.5 | 0.15 | 10 | 0.5 | 10 |
| **BI_7** | 0 | 3 | 0.15 | 50 | 0.5 | 25 |

**Figure 1.**
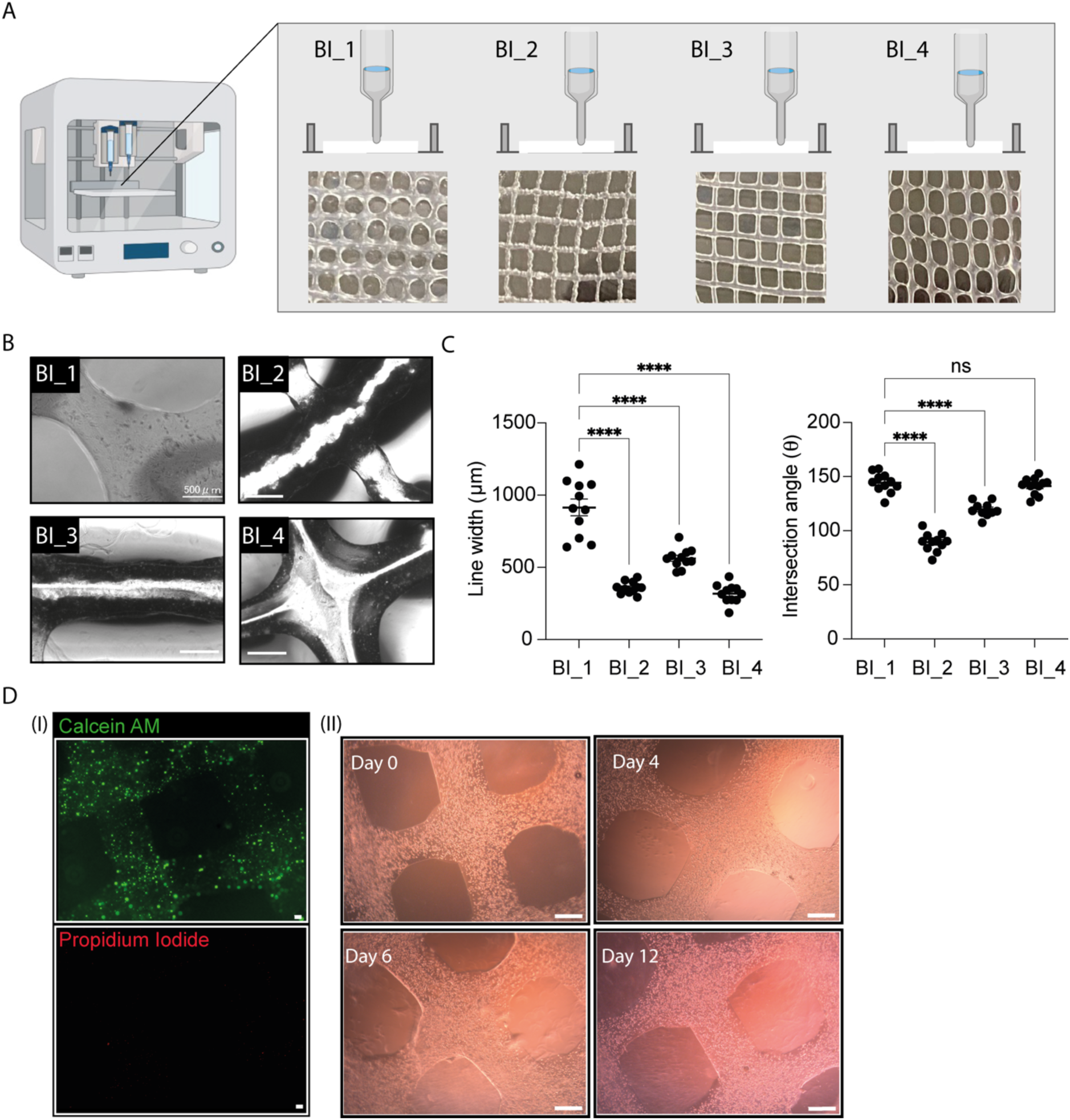
Assessment of 3D-bioprinting using alginate and gelatin bioinks. (A) Left: Image of the 3D bioprinter used for fabricating the constructs. Right: Grid-like structures printed with four different bioinks (BI_1–BI_4), demonstrating variations in print quality based on gelatin concentration. (B) Microscope images of the printed structures from each bioink. (C) Quantitative analysis of line width and intersection angles for each bioink, with statistical significance indicated (****p < 0.0001; ns: not significant). (D) (I) Live-dead assay in cells after 24 hours. Green: live cells; red: dead cells. Scale bars: 100 μm. Phase-contrast images showing cell morphology within the scaffold printed with BI_4 at days 0, 4, 6, and 12. Scale bars in (B) and (D) are 500 μm.

### 3.2. Cell differentiation in fibrinogen-infused bioinks

#### 3. 2. 1. Myoblast differentiation and spatial distribution with fibrinogen-alginate bioink

Due to the limited differentiation observed with bioinks composed solely of alginate, we hypothesized that adding a bioactive component, such as fibrinogen, could enable bioprinting at room temperature and enhance myoblast differentiation while maintaining the mechanical strength provided by alginate and gelatin. Consequently, we formulated bioink BI_5 to encapsulate C2C12 cells (**Figure 2A**). The printability of a rounded shape with a grid infill was achieved, demonstrating a mean printing fidelity index of 0.97 ± 0.08 (**Figure 2B**). We then assessed cell viability and cytoskeletal organization within the printed constructs. Calcein AM staining (**Figure 2B; (I)**) revealed high cell viability five days post-printing. Structural analysis with DAPI and F-actin staining (**Figure 2B; (II)**) showed improved cytoskeletal organization in cells encapsulated in bioink BI_5, with pronounced actin filament networks in the 3D constructs at day 5. However, shrinkage of the scaffold grids and a separation of the fibrin gel from alginate were observed after five days (data not shown). To improve the stability of the alginate-fibrinogen bioink, we incorporated 10% v/v nanofiber cellulose (NFC) and a small amount of hyaluronic acid, creating bioink BI_6. Bioprinting with C2C12 cells in a rounded shape with grid infill was comparable to the results achieved with BI_5 (**Figure 2D**). Further, Calcein AM and Hoechst 3342 staining indicated significant cell growth within the scaffolds on day five post-culturing (**Figure 2E; (I)**). Confocal scanning microscopy revealed the spatial distribution of differentiated cells, which localized mainly on the surface of the grids rather than within the scaffold core (**Figure 2F**). Quantitative analysis comparing the grid lines width of scaffolds printed with BI_5 to those printed with BI_6 demonstrated significant reduction in line width in BI_5 (**Figure 2G**).

**Figure 2.**
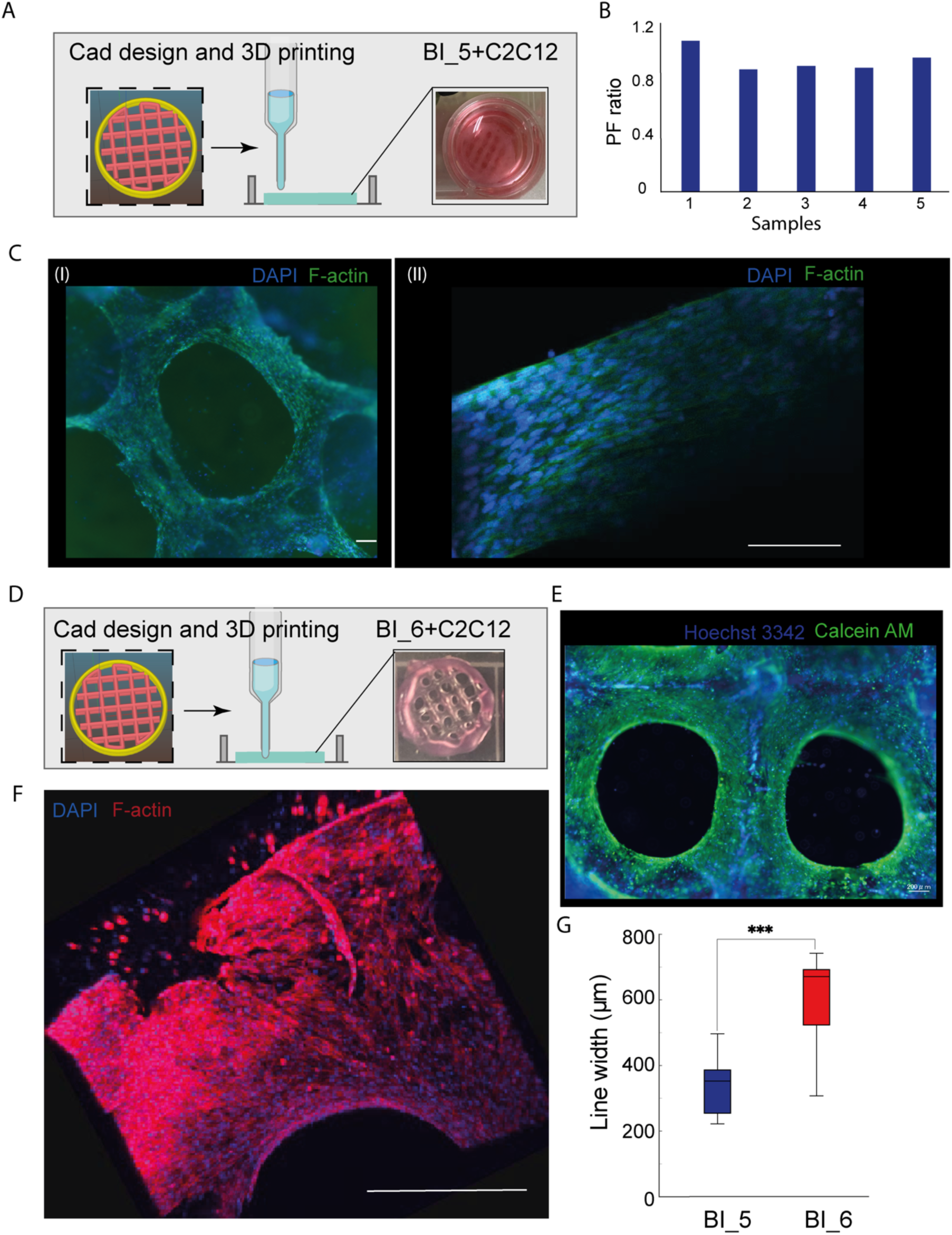
Scaffold fabrication and bioprinting using fibrinogen-alginate bioinks. (A) Scaffold design and 3D printing process: A workflow illustration from CAD design to 3D printing. The scaffolds, designed with a grid pattern (yellow and red), were printed using bioink BI_5 containing C2C12 cells. The final printed scaffold is shown in a culture dish, demonstrating its physical integrity and readiness for biological applications. (B) Graph displaying the PF ratio of five different scaffold samples, indicating consistent mechanical properties across all tested samples. Each bar represents an individual measurement, highlighting the uniformity and reliability of the 3D printing process. (C) Fluorescent images of cells on the scaffold: (I) Overview of a scaffold with cells stained using DAPI (blue) to indicate nuclei and F-actin (green) to highlight the cytoskeletal structure. (II) High-magnification confocal scanning microscopy image showing cells stained with DAPI (blue) and F-actin (green), demonstrating continuous cellular networks and cytoskeletal formations across the scaffold. (D) Scaffold design and 3D printing process: Another workflow illustration from CAD design to 3D printing. Scaffolds with a grid pattern (yellow and red) were printed using bioink BI_6 with C2C12 cells, and the final printed scaffold exhibits physical integrity and readiness for biological use. (E) Fluorescent images of cells on the scaffold: Cells stained with DAPI (blue) for nuclei and Calcein AM (green) to highlight cell viability. (F) Confocal scanning microscope images displaying the 3D structure of the scaffold with cells stained using DAPI (blue) and F-actin (green). Scale bars: 200 μm (C, I and II), 200 μm (E, I and II), and 500 μm (F). (G) Box plot showing scaffold line width produced using bioinks BI_5 and BI_6. The median for each group is represented by a black line, and the interquartile range (IQR) spans from the first quartile (Q1, 25th percentile) to the third quartile (Q3, 75th percentile) (n = 3–4). The p-values were determined using the t-test (***p < 0.001).

To develop stable 3D skeletal muscle models suitable for functional measurements under stretching and electrical stimulation, we incorporated PDMS pillars into the designs of both model 1 (a rectangular structure) and model 2 (interlocking rings) (**Figure 3A**). These PDMS pillars served as structural reinforcements during and after the 3D bioprinting process, primarily functioning as a support framework to preserve the complex geometry of the interlocking rings and the larger surface areas in the rectangular structure. The successful fabrication of these models, reinforced with PDMS pillars, allowed them to be immersed in cell culture media for biological evaluation. Despite the softness of the bioinks used, which could otherwise lead to deformation or collapse, the PDMS pillars effectively stabilized the constructs, enabling reliable cell seeding and maintaining the 3D structure’s integrity throughout the culture period. In model 1, bioink deposition resulted in some deformation of the structure, although it remained intact around the pillars. Comparatively, model 2 exhibited higher printing deposition accuracy and less structural deformation overall (**Figure 3B**). Cell viability and spatial distribution within the printed constructs were assessed using Calcein AM staining. Both models demonstrated high levels of live cells, with model 2-based scaffolds showing a higher intensity of viable cells (**Figure 3C and D; (I)**). And a higher magnification image revealed even cellular coverage and elongation along the edges of the structure (**Figure 3D; (II)**).

**Figure 3.**
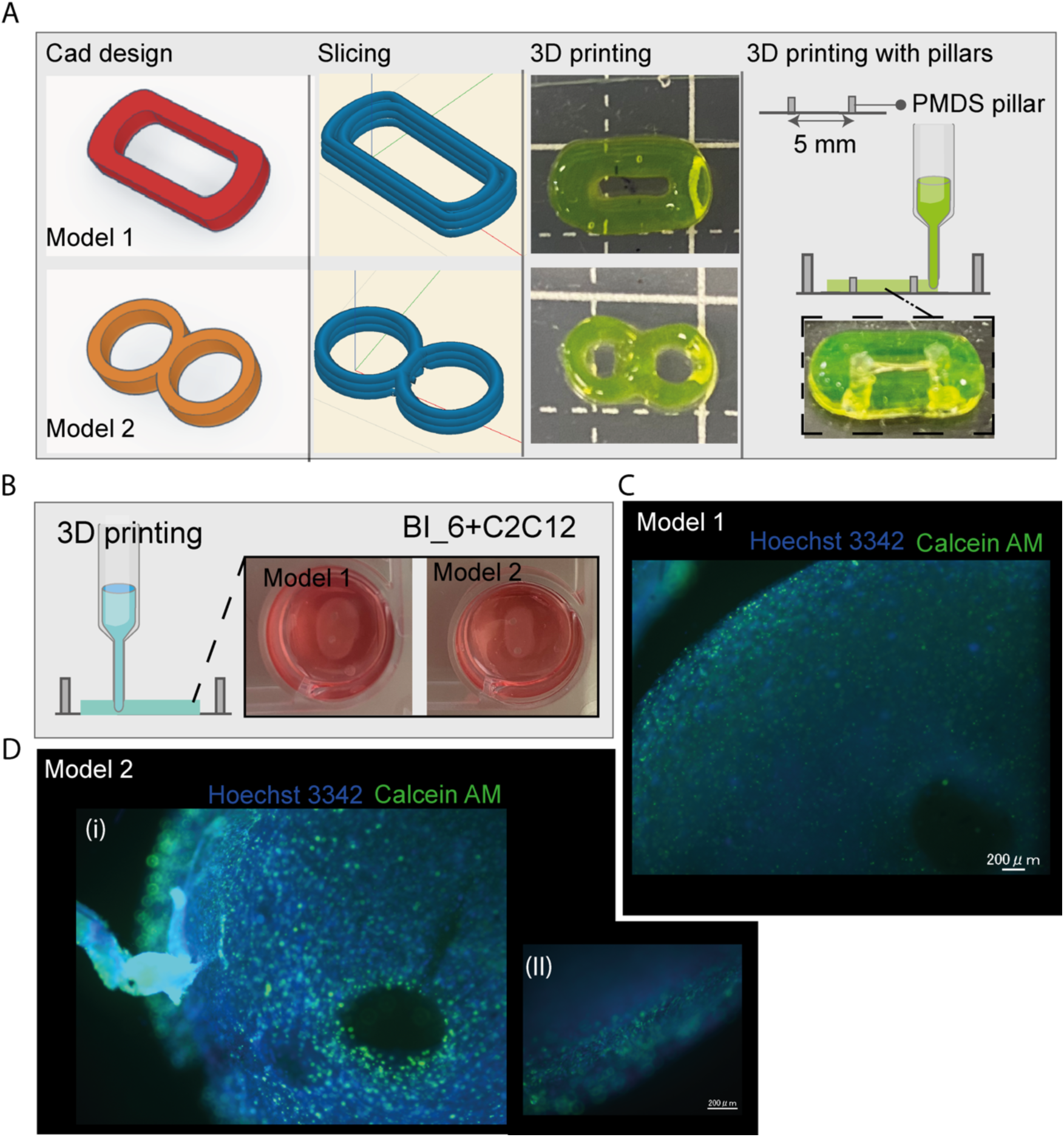
Design and characterization of 3D-bioprinted models incorporating C2C12 cells on PDMS pillars. (A) CAD designs (left) and slicing (middle left) of two models: model 1, a rectangular structure, and model 2, composed of two interlocking rings. The 3D printing process (middle right) shows the successful fabrication of both models. Models were printed on PDMS pillars for enhanced stability and structural integrity (right). (B) 3D-printed versions of model 1 and model 2 using bioink BI_5. (C) Fluorescent images of cells cultured in model 1, stained with Calcein AM (green, live cells) and Hoechst 3342 (blue, nuclei) after 6 days of culture. (D) (I) Fluorescent images of cells cultured in model 2, stained with Calcein AM (green, live cells) and Hoechst 3342 (blue, nuclei) after 6 days of culture. (II) Live-dead staining of model 2 shows the distribution of differentiated cells along the scaffold edges. Scale bars: 200 μm.

#### 3. 2. 2. Myoblast differentiation spatial distribution and functions with fibrinogen-nanofiber cellulose (NFC) bioink

We next examined the feasibility of printing constructs using a bioink composed of a high concentration of fibrinogen, gelatin, and an increased proportion of NFC as a substitute for alginate. This bioink was printable at room temperature under a pneumatic pressure of 10–15 kPa, with gelatin and NFC providing the necessary structural integrity during printing, prior to the addition of the thrombin cross-linking agent. Successful printing of interlocking ring structures (model 2) around PDMS pillars was achieved. During long-term myoblast cell culture, the rings gradually formed fused scaffolds around the PDMS pillars. As gelatin gradually released from the constructs, notable scaffold shrinkage was observed by day 16, in contrast to day 6 (**Figure 4A**). To assess myoblast differentiation and spatial distribution within the scaffolds, we conducted structural analysis via confocal scanning microscopy to visualize actin fibers in the cells. By day 6, cells had elongated significantly, showing signs of fusion, with the 3D structure promoting cell distribution throughout both the surface and core of the scaffolds (**Figure 4B; (I), (II)**). By day 16, further differentiation was evident, with clear myotube formation and alignment observed (**Figure 4C; (I), (II)**). The 3D structure remained stable, supporting well-distributed cells on the surface and within the core regions of the scaffolds, including the upper regions near the PDMS pillars and the middle regions, where aligned myotubes were apparent (**Figure 4C; (III)**) (**Supplemental video 1, 2, 3)**. To further investigate scaffold functionality, we assessed their response to electrical stimulation (ES) to evaluate contractile properties. As illustrated in **Figure 4D; I, II**, applying ES at 5V for 1 second, followed by a 4-seconds rest period, induced contractile activity in specific scaffold regions. This was confirmed by optical density analysis, which showed changes in pixel intensity (**Figure 4D; (III)**).

**Figure 4.**
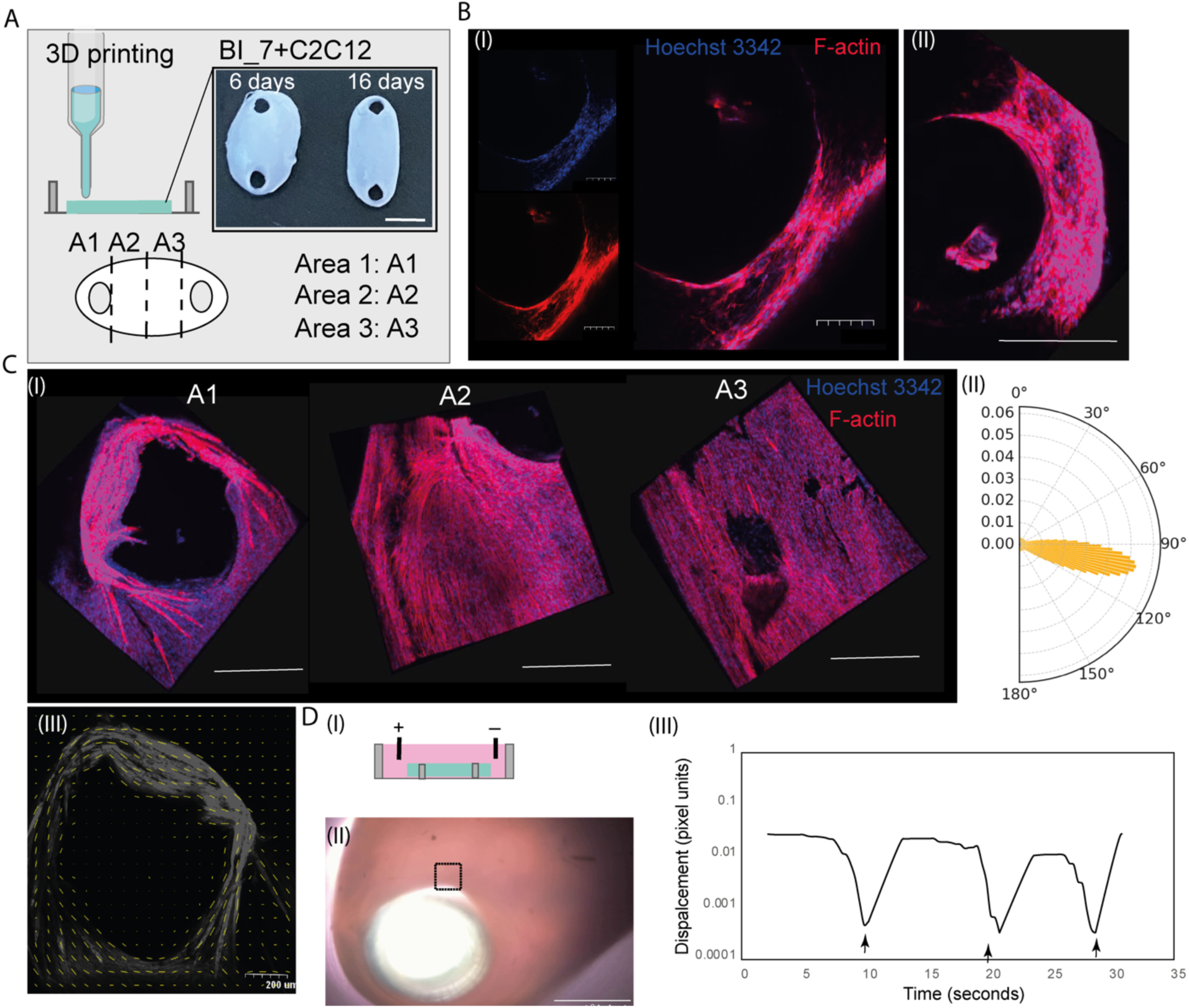
Evaluation of the effects of nanocellulose combined with fibrinogen on myoblast differentiation and spatial distribution within 3D-bioprinted scaffolds. (A) Schematic representation of the bioprinting process using bioink BI_7, illustrating the final scaffold structures at days 6 and 16. Scale bars: 2 mm. (B) Fluorescent images of scaffolds at day 6 displaying: (I) Myoblast differentiation and myotube formation, and (II) the 3D cellular structure captured using confocal scanning microscopy. Hoechst 3342 (blue) and F-actin (red) staining are used for visualization. Scale bars: 200 μm (B, I and II), and 500 μm (B, II). (C) (I) 3D visualization of myotube distribution in various scaffold regions at day 16. (II) Polar plot depicting the orientation of myotubes in regions A2 and A3. (III) Spatial analysis of cell alignment, with yellow lines indicating the direction of cell orientation on the scaffold surface. Hoechst 3342 (blue) and F-actin (red) staining are used for visualization. Scale bars: 500 μm (C). (D) (I) Illustration of the electrical stimulation setup, with carbon electrodes positioned directly on top of the scaffolds. (II) Bright-field image showing scaffolds arranged around a PDMS pillar, highlighting the marked square region used for contractile force analysis. (III) Graph depicting the displacement of cells in response to electrical stimulation, with arrows indicating the electrical triggers that cause pixel-based displacement changes in the region of interest (ROI).

## Discussion

This study highlights the critical role of bioink composition in 3D bioprinting of skeletal muscles cells, demonstrating that the interaction between bioink components significantly influences myoblast cell behaviour and scaffold performance.

Our results demonstrated that while alginate is effective in ensuring high printing fidelity, its lack of components that promote cell growth and differentiation renders it biologically inert, especially in the context of skeletal muscle cells^16^. This observation is consistent with previous findings, which have highlighted alginate’s limited ability to provide the necessary biological signalling cues for cellular activities.

To address this limitation, we incorporated fibrinogen, a soluble glycoprotein crucial in the blood clotting process. When activated by enzymes such as thrombin, fibrinogen is converted into fibrin, forming a mesh-like structure that aids in clot formation ^13^. In tissue engineering and cell culture, fibrinogen is widely utilized for its biological properties that promote cell adhesion, proliferation, and differentiation. It provides a more biologically relevant environment than inert materials like alginate, enhancing the biological signaling required for specific cell types, such as skeletal muscle cells, to grow and differentiate effectively. When we combined fibrinogen with alginate, the resulting bioink improved both cell viability and structural integrity within the printed constructs, suggesting its potential to enhance the maturation and functionality of bioprinted muscle tissues. However, the spatial distribution of cells was concentrated primarily on the surface of the scaffolds. During cell culturing, we observed a clear separation between cells embedded in the fibrin gel and the alginate fibers (data not shown), which was further confirmed by structural analysis using confocal scanning microscope. As previously mentioned, this once again highlights the inert nature of alginate, limiting cell growth within the scaffold’s core, which aligns with expectations for alginate.

To further demonstrate the capability of bioprinting in tailoring 3D *in vitro* models, we aimed to construct a scaffold that could form a muscle structure between PDMS soft pillars, a method already established for evaluating muscle contractile forces ^17,18^. Unlike previous approaches, 3D bioprinting offers the advantage of directly printing specific shapes of interest without the need for molds to constrain the cells within the ECM. We successfully printed two models: a rectangular structure (model 1) and interlocking rings (model 2). These models could be shaped around the PDMS pillars and cultured with cells for at least 16 days showcased the feasibility of bioprinting for *in vitro* modelling of skeletal muscles.

Nanocellulose refers to cellulose materials with nanoscale dimensions, typically derived from plant fibers or bacteria. Its excellent mechanical properties, biocompatibility, and high surface area make it an attractive material for tissue engineering, and 3D bioprinting. Nanocellulose is often used as a reinforcing agent in hydrogels and bioinks, as it enhances the mechanical strength and structural integrity of printed constructs while maintaining biocompatibility. Moreover, nanocellulose’s high water retention capability and nanoscale porosity mimic the natural extracellular matrix, making it particularly well-suited for scaffolds used in cell growth and tissue regeneration ^19^. Given these advantages, we incorporated NFC synthesized by acetic acid bacteria in place of alginate into our bioink, in combination with fibrinogen and gelatin. NFC along with gelatin, provided the necessary physical elasticity during the printing process, allowing the scaffold to form around the PDMS pillars while ensuring structural stability. No separation of NFC was observed after cross-linking with thrombin, indicating that the NFC remained effectively entrapped within the fibrinogen and gelatin matrix.

After incorporating myoblast cells, we investigated cell growth and differentiation. Clear signs of differentiation were observed on both day 6 and day 16, with evident formation of myotubes. By day 16, the scaffold’s shape had altered, likely due to the gradual release of the fugitive gelatin material from the structure. Interestingly, spatial analysis of the cells across the scaffold revealed successful myoblast growth and differentiation into myotubes throughout the entire scaffold. This highlights the advantage of using nanofiber cellulose as a replacement for alginate in reinforcing the fibrinogen bioink, making it more effective for bioprinting skeletal muscle cells. Additionally, the scaffolds exhibited a significant contractile response to electrical stimulation, indicating their potential use as *in vitro* models for drug response evaluation and electrophysiological analysis.

In this study, we primarily utilized murine myoblast cells. Future research should explore the use of human myoblasts and consider incorporating additional cell types, such as neurons and endothelial cells, to enhance the applicability of this approach. Furthermore, a more comprehensive analysis of the mechanical properties of NFC and their effects on cell differentiation should be addressed in further studies.

## Conclusion

In this study, we developed and characterized a series of bioinks tailored for the 3D bioprinting of skeletal muscle models. We successfully created a novel bioink formulation incorporating alginate, fibrinogen, NFC, and gelatin, demonstrating its suitability for bioprinting skeletal muscle tissues. NFC emerged as an effective alternative to alginate, improving both the mechanical properties and biological compatibility of the fibrin-based scaffolds. The bioink facilitated myoblast cell proliferation and differentiation into functional myotubes, with uniform distribution throughout the constructs **(Figure 5)**. Furthermore, the bioprinted scaffolds showed a robust contractile response to electrical stimulation, highlighting their potential as *in vitro* models for drug testing and electrophysiological analysis.

**Figure 5.**
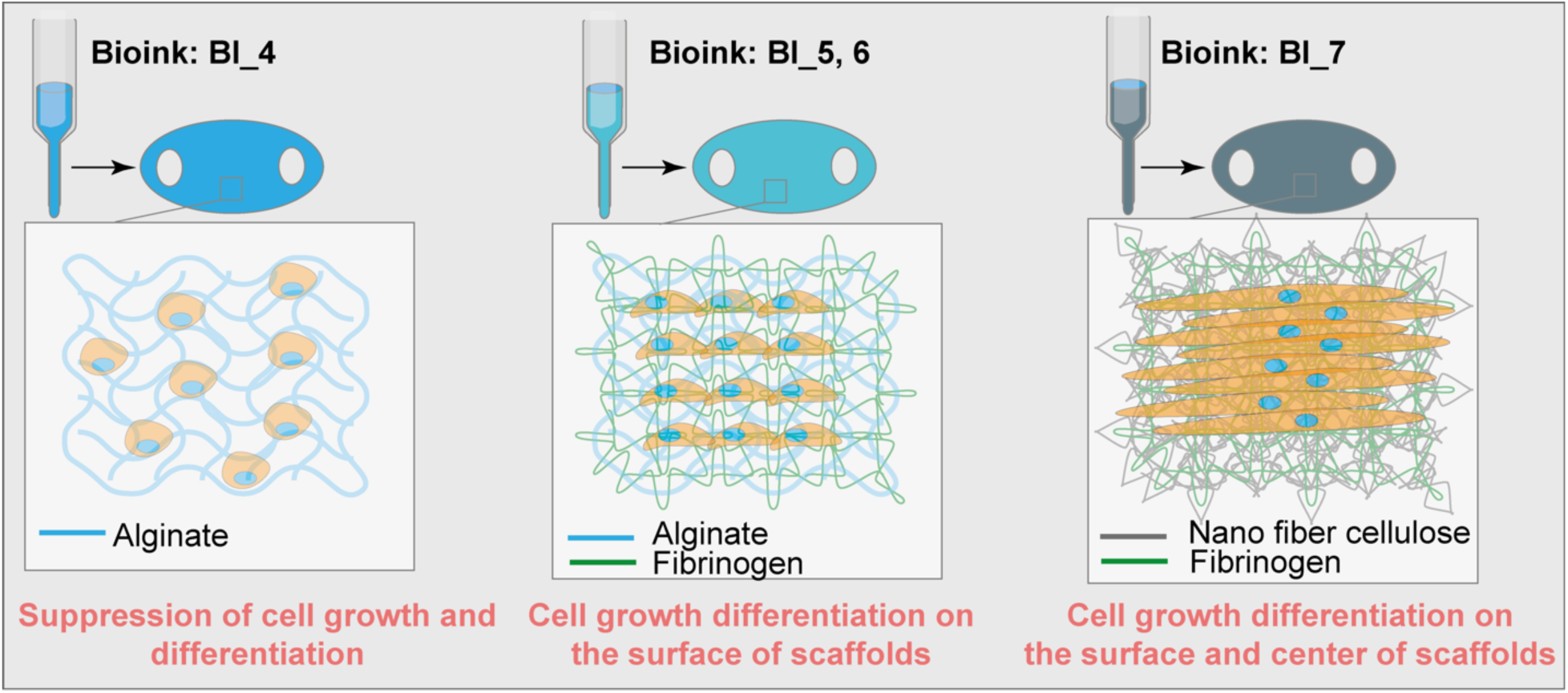
Summary of the effects of different bioink compositions on cell growth and differentiation in 3D-bioprinted scaffolds. (A) Bioink BI_4 (alginate only) suppresses cell growth and differentiation, resulting in limited cell interaction within the scaffold. (B) Bioinks BI_5 and BI_6 (alginate with fibrinogen) promote cell growth and differentiation on the surface of the scaffolds, with minimal infiltration into the scaffold structure. (C) Bioink BI_7 (nanocellulose with fibrinogen) enhances cell growth and differentiation throughout the scaffold, supporting uniform cell distribution and the formation of aligned myotubes both on the surface and within the center of the scaffold.

## Acknowledgments

This work was generously supported by the Japan Society for the Promotion of Science (JSPS, 22K14548 and 24K15712) and the Hirose Foundation, awarded to Rodi Kado Abdalkader. We also gratefully acknowledge the Ritsumeikan Global Innovation Research Organization (R-GIRO) for their support. Additionally, we thank the Biomedical Engineering Center (BMEC) at Ritsumeikan University for providing access to the confocal scanning microscopy facility. We also thanks (Nano T-Sailing, LLC. Tokushima, Japan) for providing the nanofiber cellulose material.

## Authors contribution

Rodi Kado Abdalkader was responsible for the overall research concept, project management, experimental design and execution of biological studies, data analysis and interpretation, results visualization, and manuscript writing. Kosei Yamauchi contributed to the electrical stimulation experiments, as well as data analysis and interpretation. Satoshi Konishi and Takuya Fujita provided resources and contributed to data interpretation and manuscript refinement. All authors have reviewed and approved the final version of the manuscript for publication.

## Additional information

Competing interests: All authors declare no competing financial interest.

