## Supplemental data 1 for "Development and characterization of bioinks for 3D bioprinting of in vitro skeletal muscle constructs"

**Round shape (G-code)**

```
; external perimeters extrusion width = 0.41mm
; perimeters extrusion width = 0.43mm
; infill extrusion width = 0.43mm
; solid infill extrusion width = 0.43mm
; top infill extrusion width = 0.43mm
; support material extrusion width = 0.41mm
G21 ; set units to millimeters
G90 ; use absolute coordinates
M83 ; use relative distances for extrusion
G1 Z0.400 F300.000 ; move to next layer (0)
G1 E-2.00000 F300.00000 ; retract
G1 X 0.04 Y-0.48 Z0.4 E-2.0
G1 X 0.07 Y-0.96 Z0.4 E-2.0
G1 X 0.11 Y-1.45 Z0.4 E-2.0
G1 X 0.14 Y-1.93 Z0.4 E-2.0
G1 X 0.18 Y-2.41 Z0.4 E-2.0
G1 X 0.22 Y-2.89 Z0.4 E-2.0
G1 X 0.25 Y-3.38 Z0.4 E-2.0
G1 X 0.29 Y-3.86 Z0.4 E-2.0
G1 X 0.32 Y-4.34 Z0.4 E-2.0
G1 X 0.36 Y-4.82 Z0.4 E-2.0
G1 X 0.4 Y-5.31 Z0.4 E-2.0
G1 X 0.43 Y-5.79 Z0.4 E-2.0
G1 X0.468 Y-6.272 F300.000 ; move to first perimeter point
M760 ; Activate pressure valve for PH1
G1 E2.00000 F300.00000 ; unretract
G1 X 0.85 Y-6.22 Z0.4 E1.00703
G1 X1.228 Y-6.174 E0.01406 F300.000 ; perimeter
G1 X 1.53 Y-6.1 Z0.4 E0.0127
G1 X1.827 Y-6.024 E0.01134 ; perimeter
G1 X 2.12 Y-5.92 Z0.4 E0.01133
G1 X2.408 Y-5.815 E0.01132 ; perimeter
```

G1 X 2.69 Y-5.68 Z0.4 E0.01133

G1 X2.967 Y-5.551 E0.01134 ; perimeter

G1 X 3.23 Y-5.39 Z0.4 E0.01133

G1 X3.497 Y-5.234 E0.01132 ; perimeter

G1 X 3.75 Y-5.05 Z0.4 E0.01133

G1 X3.994 Y-4.866 E0.01134 ; perimeter

G1 X 4.22 Y-4.66 Z0.4 E0.01133

G1 X4.451 Y-4.451 E0.01133 ; perimeter

G1 X 4.66 Y-4.22 Z0.4 E0.01133

G1 X4.866 Y-3.994 E0.01133 ; perimeter

G1 X 5.05 Y-3.75 Z0.4 E0.01133

G1 X5.234 Y-3.497 E0.01134 ; perimeter

G1 X 5.39 Y-3.23 Z0.4 E0.01133

G1 X5.551 Y-2.967 E0.01132 ; perimeter

G1 X 5.68 Y-2.69 Z0.4 E0.01133

G1 X5.815 Y-2.408 E0.01134 ; perimeter

G1 X 5.92 Y-2.12 Z0.4 E0.01133

G1 X6.024 Y-1.827 E0.01132 ; perimeter

G1 X 6.1 Y-1.53 Z0.4 E0.01133

G1 X6.174 Y-1.228 E0.01134 ; perimeter

G1 X 6.22 Y-0.92 Z0.4 E0.01133

G1 X6.265 Y-0.617 E0.01133 ; perimeter

G1 X 6.28 Y-0.31 Z0.4 E0.01133

G1 X6.295 Y-0.000 E0.01133 ; perimeter

G1 X 6.28 Y0.31 Z0.4 E0.01133

G1 X6.265 Y0.617 E0.01133 ; perimeter

G1 X 6.22 Y0.92 Z0.4 E0.01133

G1 X6.174 Y1.228 E0.01133 ; perimeter

G1 X 6.1 Y1.53 Z0.4 E0.01133

G1 X6.024 Y1.827 E0.01134 ; perimeter

G1 X 5.92 Y2.12 Z0.4 E0.01133

G1 X5.815 Y2.408 E0.01132 ; perimeter

G1 X 5.68 Y2.69 Z0.4 E0.01133

```
G1 X5.551 Y2.967 E0.01134 ; perimeter
G1 X 5.39 Y3.23 Z0.4 E0.01133
G1 X5.234 Y3.497 E0.01132 ; perimeter
G1 X 5.05 Y3.75 Z0.4 E0.01133
G1 X4.866 Y3.994 E0.01134 ; perimeter
G1 X 4.66 Y4.22 Z0.4 E0.01133
G1 X4.451 Y4.451 E0.01133 ; perimeter
G1 X 4.22 Y4.66 Z0.4 E0.01133
G1 X3.994 Y4.866 E0.01133 ; perimeter
G1 X 3.75 Y5.05 Z0.4 E0.01133
G1 X3.497 Y5.234 E0.01134 ; perimeter
G1 X 3.23 Y5.39 Z0.4 E0.01133
G1 X2.967 Y5.551 E0.01132 ; perimeter
G1 X 2.69 Y5.68 Z0.4 E0.01133
G1 X2.408 Y5.815 E0.01134 ; perimeter
G1 X 2.12 Y5.92 Z0.4 E0.01133
G1 X1.827 Y6.024 E0.01132 ; perimeter
G1 X 1.53 Y6.1 Z0.4 E0.01133
G1 X1.228 Y6.174 E0.01134 ; perimeter
G1 X 0.92 Y6.22 Z0.4 E0.01133
G1 X0.617 Y6.265 E0.01133 ; perimeter
G1 X 0.31 Y6.28 Z0.4 E0.01133
G1 X-0.000 Y6.295 E0.01133 ; perimeter
G1 X -0.31 Y6.28 Z0.4 E0.01133
G1 X-0.617 Y6.265 E0.01133 ; perimeter
G1 X -0.92 Y6.22 Z0.4 E0.01133
G1 X-1.228 Y6.174 E0.01133 ; perimeter
G1 X -1.53 Y6.1 Z0.4 E0.01133
G1 X-1.827 Y6.024 E0.01134 ; perimeter
G1 X -2.12 Y5.92 Z0.4 E0.01133
G1 X-2.408 Y5.815 E0.01132 ; perimeter
G1 X -2.69 Y5.68 Z0.4 E0.01133
G1 X-2.967 Y5.551 E0.01134 ; perimeter
```

```
G1 X -3.23 Y5.39 Z0.4 E0.01133
G1 X-3.497 Y5.234 E0.01132 ; perimeter
G1 X -3.75 Y5.05 Z0.4 E0.01133
G1 X-3.994 Y4.866 E0.01134 ; perimeter
G1 X -4.22 Y4.66 Z0.4 E0.01133
G1 X-4.451 Y4.451 E0.01133 ; perimeter
G1 X -4.66 Y4.22 Z0.4 E0.01133
G1 X-4.866 Y3.994 E0.01133 ; perimeter
G1 X -5.05 Y3.75 Z0.4 E0.01133
G1 X-5.234 Y3.497 E0.01134 ; perimeter
G1 X -5.39 Y3.23 Z0.4 E0.01133
G1 X-5.551 Y2.967 E0.01132 ; perimeter
G1 X -5.68 Y2.69 Z0.4 E0.01133
G1 X-5.815 Y2.408 E0.01134 ; perimeter
G1 X -5.92 Y2.12 Z0.4 E0.01133
G1 X-6.024 Y1.827 E0.01132 ; perimeter
G1 X -6.1 Y1.53 Z0.4 E0.01133
G1 X-6.174 Y1.228 E0.01134 ; perimeter
G1 X -6.22 Y0.92 Z0.4 E0.01133
G1 X-6.265 Y0.617 E0.01133 ; perimeter
G1 X -6.28 Y0.31 Z0.4 E0.01133
G1 X-6.295 Y-0.000 E0.01133 ; perimeter
G1 X -6.28 Y-0.31 Z0.4 E0.01133
G1 X-6.265 Y-0.617 E0.01133 ; perimeter
G1 X -6.22 Y-0.92 Z0.4 E0.01133
G1 X-6.174 Y-1.228 E0.01133 ; perimeter
G1 X -6.1 Y-1.53 Z0.4 E0.01133
G1 X-6.024 Y-1.827 E0.01134 ; perimeter
G1 X -5.92 Y-2.12 Z0.4 E0.01133
G1 X-5.815 Y-2.408 E0.01132 ; perimeter
G1 X -5.68 Y-2.69 Z0.4 E0.01133
G1 X-5.551 Y-2.967 E0.01134 ; perimeter
G1 X -5.39 Y-3.23 Z0.4 E0.01133
```

```
G1 X-5.234 Y-3.497 E0.01132 ; perimeter
G1 X -5.05 Y-3.75 Z0.4 E0.01133
G1 X-4.866 Y-3.994 E0.01134 ; perimeter
G1 X -4.66 Y-4.22 Z0.4 E0.01133
G1 X-4.451 Y-4.451 E0.01133 ; perimeter
G1 X -4.22 Y-4.66 Z0.4 E0.01133
G1 X-3.994 Y-4.866 E0.01133 ; perimeter
G1 X -3.75 Y-5.05 Z0.4 E0.01133
G1 X-3.497 Y-5.234 E0.01134 ; perimeter
G1 X -3.23 Y-5.39 Z0.4 E0.01133
G1 X-2.967 Y-5.551 E0.01132 ; perimeter
G1 X -2.69 Y-5.68 Z0.4 E0.01133
G1 X-2.408 Y-5.815 E0.01134 ; perimeter
G1 X -2.12 Y-5.92 Z0.4 E0.01133
G1 X-1.827 Y-6.024 E0.01132 ; perimeter
G1 X -1.53 Y-6.1 Z0.4 E0.01133
G1 X-1.228 Y-6.174 E0.01134 ; perimeter
G1 X -0.92 Y-6.22 Z0.4 E0.01133
G1 X-0.617 Y-6.265 E0.01133 ; perimeter
G1 X -0.31 Y-6.28 Z0.4 E0.01133
G1 X0.000 Y-6.295 E0.01133 ; perimeter
G1 X0.406 Y-6.275 E0.00746 ; perimeter
M761 ; Deactivate valve for PH1
G1 E-2.00000 F300.00000 ; retract
G1 X 0.85 Y-6.08 Z0.4 E-2.0
G1 X 1.3 Y-5.88 Z0.4 E-2.0
G1 X 1.74 Y-5.68 Z0.4 E-2.0
G1 X 2.19 Y-5.48 Z0.4 E-2.0
G1 X 2.64 Y-5.29 Z0.4 E-2.0
G1 X 3.08 Y-5.09 Z0.4 E-2.0
G1 X3.528 Y-4.890 F300.000 ; move to first infill point
M760 ; Activate pressure valve for PH1
G1 E2.00000 F300.00000 ; unretract
```

```
G1 X 3.06 Y-4.89 Z0.4 E1.87584
G1 X 2.59 Y-4.89 Z0.4 E1.75168
G1 X 2.12 Y-4.89 Z0.4 E1.62752
G1 X 1.65 Y-4.89 Z0.4 E1.50337
G1 X 1.18 Y-4.89 Z0.4 E1.37921
G1 X 0.71 Y-4.89 Z0.4 E1.25505
G1 X 0.24 Y-4.89 Z0.4 E1.13089
G1 X -0.24 Y-4.89 Z0.4 E1.00673
G1 X -0.71 Y-4.89 Z0.4 E0.88257
G1 X -1.18 Y-4.89 Z0.4 E0.75841
G1 X -1.65 Y-4.89 Z0.4 E0.63425
G1 X -2.12 Y-4.89 Z0.4 E0.5101
G1 X -2.59 Y-4.89 Z0.4 E0.38594
G1 X -3.06 Y-4.89 Z0.4 E0.26178
G1 X-3.528 Y-4.890 E0.13762 F300.000 ; infill
G1 X -3.83 Y-4.55 Z0.4 E0.12353
G1 X -4.13 Y-4.21 Z0.4 E0.10944
G1 X -4.44 Y-3.88 Z0.4 E0.09535
G1 X -4.74 Y-3.54 Z0.4 E0.08127
G1 X -5.04 Y-3.2 Z0.4 E0.06718
G1 X-5.344 Y-2.863 E0.05309 ; infill
G1 X -4.86 Y-2.86 Z0.4 E0.06015
G1 X -4.37 Y-2.86 Z0.4 E0.06722
G1 X -3.89 Y-2.86 Z0.4 E0.07428
G1 X -3.4 Y-2.86 Z0.4 E0.08134
G1 X -2.91 Y-2.86 Z0.4 E0.08841
G1 X -2.43 Y-2.86 Z0.4 E0.09547
G1 X -1.94 Y-2.86 Z0.4 E0.10253
G1 X -1.46 Y-2.86 Z0.4 E0.1096
G1 X -0.97 Y-2.86 Z0.4 E0.11666
G1 X -0.49 Y-2.86 Z0.4 E0.12372
G1 X 0.0 Y-2.86 Z0.4 E0.13078
G1 X 0.49 Y-2.86 Z0.4 E0.13785
```

```
G1 X 0.97 Y-2.86 Z0.4 E0.14491
G1 X 1.46 Y-2.86 Z0.4 E0.15197
G1 X 1.94 Y-2.86 Z0.4 E0.15904
G1 X 2.43 Y-2.86 Z0.4 E0.1661
G1 X 2.91 Y-2.86 Z0.4 E0.17316
G1 X 3.4 Y-2.86 Z0.4 E0.18023
G1 X 3.89 Y-2.86 Z0.4 E0.18729
G1 X 4.37 Y-2.86 Z0.4 E0.19435
G1 X 4.86 Y-2.86 Z0.4 E0.20142
G1 X5.344 Y-2.863 E0.20848 ; infill
G1 X 5.48 Y-2.46 Z0.4 E0.17511
G1 X 5.61 Y-2.05 Z0.4 E0.14174
G1 X 5.75 Y-1.65 Z0.4 E0.10837
G1 X 5.88 Y-1.24 Z0.4 E0.075
G1 X6.013 Y-0.836 E0.04163 ; infill
G1 X 5.53 Y-0.84 Z0.4 E0.04935
G1 X 5.05 Y-0.84 Z0.4 E0.05706
G1 X 4.57 Y-0.84 Z0.4 E0.06478
G1 X 4.09 Y-0.84 Z0.4 E0.07249
G1 X 3.61 Y-0.84 Z0.4 E0.08021
G1 X 3.13 Y-0.84 Z0.4 E0.08793
G1 X 2.65 Y-0.84 Z0.4 E0.09564
G1 X 2.16 Y-0.84 Z0.4 E0.10336
G1 X 1.68 Y-0.84 Z0.4 E0.11107
G1 X 1.2 Y-0.84 Z0.4 E0.11879
G1 X 0.72 Y-0.84 Z0.4 E0.12651
G1 X 0.24 Y-0.84 Z0.4 E0.13422
G1 X -0.24 Y-0.84 Z0.4 E0.14194
G1 X -0.72 Y-0.84 Z0.4 E0.14965
G1 X -1.2 Y-0.84 Z0.4 E0.15737
G1 X -1.68 Y-0.84 Z0.4 E0.16509
G1 X -2.16 Y-0.84 Z0.4 E0.1728
G1 X -2.65 Y-0.84 Z0.4 E0.18052
```

```
G1 X -3.13 Y-0.84 Z0.4 E0.18823
G1 X -3.61 Y-0.84 Z0.4 E0.19595
G1 X -4.09 Y-0.84 Z0.4 E0.20367
G1 X -4.57 Y-0.84 Z0.4 E0.21138
G1 X -5.05 Y-0.84 Z0.4 E0.2191
G1 X -5.53 Y-0.84 Z0.4 E0.22681
G1 X-6.013 Y-0.836 E0.23453 ; infill
G1 X -6.0 Y-0.43 Z0.4 E0.19554
G1 X -5.99 Y-0.02 Z0.4 E0.15654
G1 X -5.98 Y0.38 Z0.4 E0.11755
G1 X -5.97 Y0.79 Z0.4 E0.07855
G1 X-5.957 Y1.192 E0.03956 ; infill
G1 X -5.46 Y1.19 Z0.4 E0.04759
G1 X -4.96 Y1.19 Z0.4 E0.05563
G1 X -4.47 Y1.19 Z0.4 E0.06366
G1 X -3.97 Y1.19 Z0.4 E0.07169
G1 X -3.47 Y1.19 Z0.4 E0.07973
G1 X -2.98 Y1.19 Z0.4 E0.08776
G1 X -2.48 Y1.19 Z0.4 E0.09579
G1 X -1.99 Y1.19 Z0.4 E0.10383
G1 X -1.49 Y1.19 Z0.4 E0.11186
G1 X -0.99 Y1.19 Z0.4 E0.11989
G1 X -0.5 Y1.19 Z0.4 E0.12793
G1 X 0.0 Y1.19 Z0.4 E0.13596
G1 X 0.5 Y1.19 Z0.4 E0.14399
G1 X 0.99 Y1.19 Z0.4 E0.15203
G1 X 1.49 Y1.19 Z0.4 E0.16006
G1 X 1.99 Y1.19 Z0.4 E0.16809
G1 X 2.48 Y1.19 Z0.4 E0.17613
G1 X 2.98 Y1.19 Z0.4 E0.18416
G1 X 3.47 Y1.19 Z0.4 E0.19219
G1 X 3.97 Y1.19 Z0.4 E0.20023
G1 X 4.47 Y1.19 Z0.4 E0.20826
```

```
G1 X 4.96 Y1.19 Z0.4 E0.21629
G1 X 5.46 Y1.19 Z0.4 E0.22433
G1 X5.957 Y1.192 E0.23236 ; infill
G1 X 5.79 Y1.6 Z0.4 E0.19443
G1 X 5.63 Y2.0 Z0.4 E0.15649
G1 X 5.46 Y2.41 Z0.4 E0.11856
G1 X 5.3 Y2.81 Z0.4 E0.08062
G1 X5.131 Y3.219 E0.04269 ; infill
G1 X 4.64 Y3.22 Z0.4 E0.05019
G1 X 4.15 Y3.22 Z0.4 E0.05769
G1 X 3.67 Y3.22 Z0.4 E0.06519
G1 X 3.18 Y3.22 Z0.4 E0.07269
G1 X 2.69 Y3.22 Z0.4 E0.08019
G1 X 2.2 Y3.22 Z0.4 E0.08768
G1 X 1.71 Y3.22 Z0.4 E0.09518
G1 X 1.22 Y3.22 Z0.4 E0.10268
G1 X 0.73 Y3.22 Z0.4 E0.11018
G1 X 0.24 Y3.22 Z0.4 E0.11768
G1 X -0.24 Y3.22 Z0.4 E0.12518
G1 X -0.73 Y3.22 Z0.4 E0.13268
G1 X -1.22 Y3.22 Z0.4 E0.14018
G1 X -1.71 Y3.22 Z0.4 E0.14768
G1 X -2.2 Y3.22 Z0.4 E0.15518
G1 X -2.69 Y3.22 Z0.4 E0.16267
G1 X -3.18 Y3.22 Z0.4 E0.17017
G1 X -3.67 Y3.22 Z0.4 E0.17767
G1 X -4.15 Y3.22 Z0.4 E0.18517
G1 X -4.64 Y3.22 Z0.4 E0.19267
G1 X-5.131 Y3.219 E0.20017 ; infill
G1 X -4.77 Y3.56 Z0.4 E0.17647
G1 X -4.41 Y3.9 Z0.4 E0.15277
G1 X -4.04 Y4.23 Z0.4 E0.12907
G1 X -3.68 Y4.57 Z0.4 E0.10538
```

```
G1 X -3.32 Y4.91 Z0.4 E0.08168
G1 X-2.957 Y5.247 E0.05798 ; infill
G1 X -2.46 Y5.25 Z0.4 E0.06276
G1 X -1.97 Y5.25 Z0.4 E0.06754
G1 X -1.48 Y5.25 Z0.4 E0.07232
G1 X -0.99 Y5.25 Z0.4 E0.0771
G1 X -0.49 Y5.25 Z0.4 E0.08188
G1 X 0.0 Y5.25 Z0.4 E0.08666
G1 X 0.49 Y5.25 Z0.4 E0.09145
G1 X 0.99 Y5.25 Z0.4 E0.09623
G1 X 1.48 Y5.25 Z0.4 E0.10101
G1 X 1.97 Y5.25 Z0.4 E0.10579
G1 X 2.46 Y5.25 Z0.4 E0.11057
G1 X2.957 Y5.247 E0.11535 ; infill
M761 ; Deactivate valve for PH1
G1 Z0.800 F300.000 ; move to next layer (1)
G1 E-2.00000 F300.00000 ; retract
G1 X 2.86 Y4.77 Z0.8 E-2.0
G1 X 2.76 Y4.29 Z0.8 E-2.0
G1 X 2.66 Y3.81 Z0.8 E-2.0
G1 X 2.57 Y3.33 Z0.8 E-2.0
G1 X 2.47 Y2.85 Z0.8 E-2.0
G1 X 2.37 Y2.37 Z0.8 E-2.0
G1 X 2.27 Y1.89 Z0.8 E-2.0
G1 X 2.18 Y1.41 Z0.8 E-2.0
G1 X 2.08 Y0.93 Z0.8 E-2.0
G1 X 1.98 Y0.45 Z0.8 E-2.0
G1 X 1.88 Y-0.03 Z0.8 E-2.0
G1 X 1.79 Y-0.51 Z0.8 E-2.0
G1 X 1.69 Y-0.99 Z0.8 E-2.0
G1 X 1.59 Y-1.47 Z0.8 E-2.0
G1 X 1.49 Y-1.95 Z0.8 E-2.0
G1 X 1.4 Y-2.43 Z0.8 E-2.0
```

G1 X 1.3 Y-2.91 Z0.8 E-2.0

G1 X 1.2 Y-3.39 Z0.8 E-2.0

G1 X 1.1 Y-3.87 Z0.8 E-2.0

G1 X 1.01 Y-4.35 Z0.8 E-2.0

G1 X 0.91 Y-4.83 Z0.8 E-2.0

G1 X 0.81 Y-5.31 Z0.8 E-2.0

G1 X 0.71 Y-5.79 Z0.8 E-2.0

G1 X0.617 Y-6.265 F300.000 ; move to first perimeter point

M760 ; Activate pressure valve for PH1

G1 E2.00000 F300.00000 ; unretract

G1 X 0.92 Y-6.22 Z0.8 E1.00567

G1 X1.228 Y-6.174 E0.01133 F300.000 ; perimeter

G1 X 1.53 Y-6.1 Z0.8 E0.01133

G1 X1.827 Y-6.024 E0.01134 ; perimeter

G1 X 2.12 Y-5.92 Z0.8 E0.01133

G1 X2.408 Y-5.815 E0.01132 ; perimeter

G1 X 2.69 Y-5.68 Z0.8 E0.01133

G1 X2.967 Y-5.551 E0.01134 ; perimeter

G1 X 3.23 Y-5.39 Z0.8 E0.01133

G1 X3.497 Y-5.234 E0.01132 ; perimeter

G1 X 3.75 Y-5.05 Z0.8 E0.01133

G1 X3.994 Y-4.866 E0.01134 ; perimeter

G1 X 4.22 Y-4.66 Z0.8 E0.01133

G1 X4.451 Y-4.451 E0.01133 ; perimeter

G1 X 4.66 Y-4.22 Z0.8 E0.01133

G1 X4.866 Y-3.994 E0.01133 ; perimeter

G1 X 5.05 Y-3.75 Z0.8 E0.01133

G1 X5.234 Y-3.497 E0.01134 ; perimeter

G1 X 5.39 Y-3.23 Z0.8 E0.01133

G1 X5.551 Y-2.967 E0.01132 ; perimeter

G1 X 5.68 Y-2.69 Z0.8 E0.01133

G1 X5.815 Y-2.408 E0.01134 ; perimeter

G1 X 5.92 Y-2.12 Z0.8 E0.01133

```
G1 X6.024 Y-1.827 E0.01132 ; perimeter
G1 X 6.1 Y-1.53 Z0.8 E0.01133
G1 X6.174 Y-1.228 E0.01134 ; perimeter
G1 X 6.22 Y-0.92 Z0.8 E0.01133
G1 X6.265 Y-0.617 E0.01133 ; perimeter
G1 X 6.28 Y-0.31 Z0.8 E0.01133
G1 X6.295 Y-0.000 E0.01133 ; perimeter
G1 X 6.28 Y0.31 Z0.8 E0.01133
G1 X6.265 Y0.617 E0.01133 ; perimeter
G1 X 6.22 Y0.92 Z0.8 E0.01133
G1 X6.174 Y1.228 E0.01133 ; perimeter
G1 X 6.1 Y1.53 Z0.8 E0.01133
G1 X6.024 Y1.827 E0.01134 ; perimeter
G1 X 5.92 Y2.12 Z0.8 E0.01133
G1 X5.815 Y2.408 E0.01132 ; perimeter
G1 X 5.68 Y2.69 Z0.8 E0.01133
G1 X5.551 Y2.967 E0.01134 ; perimeter
G1 X 5.39 Y3.23 Z0.8 E0.01133
G1 X5.234 Y3.497 E0.01132 ; perimeter
G1 X 5.05 Y3.75 Z0.8 E0.01133
G1 X4.866 Y3.994 E0.01134 ; perimeter
G1 X 4.66 Y4.22 Z0.8 E0.01133
G1 X4.451 Y4.451 E0.01133 ; perimeter
G1 X 4.22 Y4.66 Z0.8 E0.01133
G1 X3.994 Y4.866 E0.01133 ; perimeter
G1 X 3.75 Y5.05 Z0.8 E0.01133
G1 X3.497 Y5.234 E0.01134 ; perimeter
G1 X 3.23 Y5.39 Z0.8 E0.01133
G1 X2.967 Y5.551 E0.01132 ; perimeter
G1 X 2.69 Y5.68 Z0.8 E0.01133
G1 X2.408 Y5.815 E0.01134 ; perimeter
G1 X 2.12 Y5.92 Z0.8 E0.01133
G1 X1.827 Y6.024 E0.01132 ; perimeter
```

```
G1 X 1.53 Y6.1 Z0.8 E0.01133
G1 X1.228 Y6.174 E0.01134 ; perimeter
G1 X 0.92 Y6.22 Z0.8 E0.01133
G1 X0.617 Y6.265 E0.01133 ; perimeter
G1 X 0.31 Y6.28 Z0.8 E0.01133
G1 X-0.000 Y6.295 E0.01133 ; perimeter
G1 X -0.31 Y6.28 Z0.8 E0.01133
G1 X-0.617 Y6.265 E0.01133 ; perimeter
G1 X -0.92 Y6.22 Z0.8 E0.01133
G1 X-1.228 Y6.174 E0.01133 ; perimeter
G1 X -1.53 Y6.1 Z0.8 E0.01133
G1 X-1.827 Y6.024 E0.01134 ; perimeter
G1 X -2.12 Y5.92 Z0.8 E0.01133
G1 X-2.408 Y5.815 E0.01132 ; perimeter
G1 X -2.69 Y5.68 Z0.8 E0.01133
G1 X-2.967 Y5.551 E0.01134 ; perimeter
G1 X -3.23 Y5.39 Z0.8 E0.01133
G1 X-3.497 Y5.234 E0.01132 ; perimeter
G1 X -3.75 Y5.05 Z0.8 E0.01133
G1 X-3.994 Y4.866 E0.01134 ; perimeter
G1 X -4.22 Y4.66 Z0.8 E0.01133
G1 X-4.451 Y4.451 E0.01133 ; perimeter
G1 X -4.66 Y4.22 Z0.8 E0.01133
G1 X-4.866 Y3.994 E0.01133 ; perimeter
G1 X -5.05 Y3.75 Z0.8 E0.01133
G1 X-5.234 Y3.497 E0.01134 ; perimeter
G1 X -5.39 Y3.23 Z0.8 E0.01133
G1 X-5.551 Y2.967 E0.01132 ; perimeter
G1 X -5.68 Y2.69 Z0.8 E0.01133
G1 X-5.815 Y2.408 E0.01134 ; perimeter
G1 X -5.92 Y2.12 Z0.8 E0.01133
G1 X-6.024 Y1.827 E0.01132 ; perimeter
G1 X -6.1 Y1.53 Z0.8 E0.01133
```

```
G1 X-6.174 Y1.228 E0.01134 ; perimeter
G1 X -6.22 Y0.92 Z0.8 E0.01133
G1 X-6.265 Y0.617 E0.01133 ; perimeter
G1 X -6.28 Y0.31 Z0.8 E0.01133
G1 X-6.295 Y-0.000 E0.01133 ; perimeter
G1 X -6.28 Y-0.31 Z0.8 E0.01133
G1 X-6.265 Y-0.617 E0.01133 ; perimeter
G1 X -6.22 Y-0.92 Z0.8 E0.01133
G1 X-6.174 Y-1.228 E0.01133 ; perimeter
G1 X -6.1 Y-1.53 Z0.8 E0.01133
G1 X-6.024 Y-1.827 E0.01134 ; perimeter
G1 X -5.92 Y-2.12 Z0.8 E0.01133
G1 X-5.815 Y-2.408 E0.01132 ; perimeter
G1 X -5.68 Y-2.69 Z0.8 E0.01133
G1 X-5.551 Y-2.967 E0.01134 ; perimeter
G1 X -5.39 Y-3.23 Z0.8 E0.01133
G1 X-5.234 Y-3.497 E0.01132 ; perimeter
G1 X -5.05 Y-3.75 Z0.8 E0.01133
G1 X-4.866 Y-3.994 E0.01134 ; perimeter
G1 X -4.66 Y-4.22 Z0.8 E0.01133
G1 X-4.451 Y-4.451 E0.01133 ; perimeter
G1 X -4.22 Y-4.66 Z0.8 E0.01133
G1 X-3.994 Y-4.866 E0.01133 ; perimeter
G1 X -3.75 Y-5.05 Z0.8 E0.01133
G1 X-3.497 Y-5.234 E0.01134 ; perimeter
G1 X -3.23 Y-5.39 Z0.8 E0.01133
G1 X-2.967 Y-5.551 E0.01132 ; perimeter
G1 X -2.69 Y-5.68 Z0.8 E0.01133
G1 X-2.408 Y-5.815 E0.01134 ; perimeter
G1 X -2.12 Y-5.92 Z0.8 E0.01133
G1 X-1.827 Y-6.024 E0.01132 ; perimeter
G1 X -1.53 Y-6.1 Z0.8 E0.01133
G1 X-1.228 Y-6.174 E0.01134 ; perimeter
```

```
G1 X -0.92 Y-6.22 Z0.8 E0.01132
G1 X-0.618 Y-6.265 E0.01130 ; perimeter
G1 X -0.25 Y-6.28 Z0.8 E0.01233
G1 X0.110 Y-6.289 E0.01337 ; perimeter
G1 X0.555 Y-6.268 E0.00818 ; perimeter
M761 ; Deactivate valve for PH1
G1 E-2.00000 F300.00000 ; retract
G1 X 0.75 Y-5.82 Z0.8 E-2.0
G1 X 0.95 Y-5.38 Z0.8 E-2.0
G1 X 1.15 Y-4.93 Z0.8 E-2.0
G1 X 1.34 Y-4.49 Z0.8 E-2.0
G1 X 1.54 Y-4.04 Z0.8 E-2.0
G1 X 1.74 Y-3.6 Z0.8 E-2.0
G1 X 1.93 Y-3.15 Z0.8 E-2.0
G1 X 2.13 Y-2.71 Z0.8 E-2.0
G1 X 2.33 Y-2.26 Z0.8 E-2.0
G1 X 2.53 Y-1.82 Z0.8 E-2.0
G1 X 2.72 Y-1.37 Z0.8 E-2.0
G1 X 2.92 Y-0.92 Z0.8 E-2.0
G1 X 3.12 Y-0.48 Z0.8 E-2.0
G1 X 3.31 Y-0.03 Z0.8 E-2.0
G1 X 3.51 Y0.41 Z0.8 E-2.0
G1 X 3.71 Y0.86 Z0.8 E-2.0
G1 X 3.9 Y1.3 Z0.8 E-2.0
G1 X 4.1 Y1.75 Z0.8 E-2.0
G1 X 4.3 Y2.19 Z0.8 E-2.0
G1 X 4.5 Y2.64 Z0.8 E-2.0
G1 X 4.69 Y3.08 Z0.8 E-2.0
G1 X4.890 Y3.528 F300.000 ; move to first infill point
M760 ; Activate pressure valve for PH1
G1 E2.00000 F300.00000 ; unretract
G1 X 4.89 Y3.06 Z0.8 E1.87584
G1 X 4.89 Y2.59 Z0.8 E1.75168
```

```
G1 X 4.89 Y2.12 Z0.8 E1.62752
G1 X 4.89 Y1.65 Z0.8 E1.50337
G1 X 4.89 Y1.18 Z0.8 E1.37921
G1 X 4.89 Y0.71 Z0.8 E1.25505
G1 X 4.89 Y0.24 Z0.8 E1.13089
G1 X 4.89 Y-0.24 Z0.8 E1.00673
G1 X 4.89 Y-0.71 Z0.8 E0.88257
G1 X 4.89 Y-1.18 Z0.8 E0.75841
G1 X 4.89 Y-1.65 Z0.8 E0.63425
G1 X 4.89 Y-2.12 Z0.8 E0.5101
G1 X 4.89 Y-2.59 Z0.8 E0.38594
G1 X 4.89 Y-3.06 Z0.8 E0.26178
G1 X4.890 Y-3.528 E0.13762 F300.000 ; infill
G1 X 4.55 Y-3.83 Z0.8 E0.12353
G1 X 4.21 Y-4.13 Z0.8 E0.10944
G1 X 3.88 Y-4.44 Z0.8 E0.09535
G1 X 3.54 Y-4.74 Z0.8 E0.08127
G1 X 3.2 Y-5.04 Z0.8 E0.06718
G1 X2.863 Y-5.344 E0.05309 ; infill
G1 X 2.86 Y-4.86 Z0.8 E0.06015
G1 X 2.86 Y-4.37 Z0.8 E0.06722
G1 X 2.86 Y-3.89 Z0.8 E0.07428
G1 X 2.86 Y-3.4 Z0.8 E0.08134
G1 X 2.86 Y-2.91 Z0.8 E0.08841
G1 X 2.86 Y-2.43 Z0.8 E0.09547
G1 X 2.86 Y-1.94 Z0.8 E0.10253
G1 X 2.86 Y-1.46 Z0.8 E0.1096
G1 X 2.86 Y-0.97 Z0.8 E0.11666
G1 X 2.86 Y-0.49 Z0.8 E0.12372
G1 X 2.86 Y0.0 Z0.8 E0.13078
G1 X 2.86 Y0.49 Z0.8 E0.13785
G1 X 2.86 Y0.97 Z0.8 E0.14491
G1 X 2.86 Y1.46 Z0.8 E0.15197
```

G1 X 2.86 Y1.94 Z0.8 E0.15904

G1 X 2.86 Y2.43 Z0.8 E0.1661

G1 X 2.86 Y2.91 Z0.8 E0.17316

G1 X 2.86 Y3.4 Z0.8 E0.18023

G1 X 2.86 Y3.89 Z0.8 E0.18729

G1 X 2.86 Y4.37 Z0.8 E0.19435

G1 X 2.86 Y4.86 Z0.8 E0.20142

G1 X2.863 Y5.344 E0.20848 ; infill

G1 X 2.46 Y5.48 Z0.8 E0.17511

G1 X 2.05 Y5.61 Z0.8 E0.14174

G1 X 1.65 Y5.75 Z0.8 E0.10837

G1 X 1.24 Y5.88 Z0.8 E0.075

G1 X0.836 Y6.013 E0.04163 ; infill

G1 X 0.84 Y5.53 Z0.8 E0.04935

G1 X 0.84 Y5.05 Z0.8 E0.05706

G1 X 0.84 Y4.57 Z0.8 E0.06478

G1 X 0.84 Y4.09 Z0.8 E0.07249

G1 X 0.84 Y3.61 Z0.8 E0.08021

G1 X 0.84 Y3.13 Z0.8 E0.08793

G1 X 0.84 Y2.65 Z0.8 E0.09564

G1 X 0.84 Y2.16 Z0.8 E0.10336

G1 X 0.84 Y1.68 Z0.8 E0.11107

G1 X 0.84 Y1.2 Z0.8 E0.11879

G1 X 0.84 Y0.72 Z0.8 E0.12651

G1 X 0.84 Y0.24 Z0.8 E0.13422

G1 X 0.84 Y-0.24 Z0.8 E0.14194

G1 X 0.84 Y-0.72 Z0.8 E0.14965

G1 X 0.84 Y-1.2 Z0.8 E0.15737

G1 X 0.84 Y-1.68 Z0.8 E0.16509

G1 X 0.84 Y-2.16 Z0.8 E0.1728

G1 X 0.84 Y-2.65 Z0.8 E0.18052

G1 X 0.84 Y-3.13 Z0.8 E0.18823

G1 X 0.84 Y-3.61 Z0.8 E0.19595

```
G1 X 0.84 Y-4.09 Z0.8 E0.20367
G1 X 0.84 Y-4.57 Z0.8 E0.21138
G1 X 0.84 Y-5.05 Z0.8 E0.2191
G1 X 0.84 Y-5.53 Z0.8 E0.22681
G1 X0.836 Y-6.013 E0.23453 ; infill
G1 X 0.43 Y-6.0 Z0.8 E0.19554
G1 X 0.02 Y-5.99 Z0.8 E0.15654
G1 X -0.38 Y-5.98 Z0.8 E0.11755
G1 X -0.79 Y-5.97 Z0.8 E0.07855
G1 X-1.192 Y-5.957 E0.03956 ; infill
G1 X -1.19 Y-5.46 Z0.8 E0.04759
G1 X -1.19 Y-4.96 Z0.8 E0.05563
G1 X -1.19 Y-4.47 Z0.8 E0.06366
G1 X -1.19 Y-3.97 Z0.8 E0.07169
G1 X -1.19 Y-3.47 Z0.8 E0.07973
G1 X -1.19 Y-2.98 Z0.8 E0.08776
G1 X -1.19 Y-2.48 Z0.8 E0.09579
G1 X -1.19 Y-1.99 Z0.8 E0.10383
G1 X -1.19 Y-1.49 Z0.8 E0.11186
G1 X -1.19 Y-0.99 Z0.8 E0.11989
G1 X -1.19 Y-0.5 Z0.8 E0.12793
G1 X -1.19 Y0.0 Z0.8 E0.13596
G1 X -1.19 Y0.5 Z0.8 E0.14399
G1 X -1.19 Y0.99 Z0.8 E0.15203
G1 X -1.19 Y1.49 Z0.8 E0.16006
G1 X -1.19 Y1.99 Z0.8 E0.16809
G1 X -1.19 Y2.48 Z0.8 E0.17613
G1 X -1.19 Y2.98 Z0.8 E0.18416
G1 X -1.19 Y3.47 Z0.8 E0.19219
G1 X -1.19 Y3.97 Z0.8 E0.20023
G1 X -1.19 Y4.47 Z0.8 E0.20826
G1 X -1.19 Y4.96 Z0.8 E0.21629
G1 X -1.19 Y5.46 Z0.8 E0.22433
```

```
G1 X-1.192 Y5.957 E0.23236 ; infill
G1 X -1.6 Y5.79 Z0.8 E0.19443
G1 X -2.0 Y5.63 Z0.8 E0.15649
G1 X -2.41 Y5.46 Z0.8 E0.11856
G1 X -2.81 Y5.3 Z0.8 E0.08062
G1 X-3.219 Y5.131 E0.04269 ; infill
G1 X -3.22 Y4.64 Z0.8 E0.05019
G1 X -3.22 Y4.15 Z0.8 E0.05769
G1 X -3.22 Y3.66 Z0.8 E0.06519
G1 X -3.22 Y3.18 Z0.8 E0.07269
G1 X -3.22 Y2.69 Z0.8 E0.08019
G1 X -3.22 Y2.2 Z0.8 E0.08768
G1 X -3.22 Y1.71 Z0.8 E0.09518
G1 X -3.22 Y1.22 Z0.8 E0.10268
G1 X -3.22 Y0.73 Z0.8 E0.11018
G1 X -3.22 Y0.24 Z0.8 E0.11768
G1 X -3.22 Y-0.24 Z0.8 E0.12518
G1 X -3.22 Y-0.73 Z0.8 E0.13268
G1 X -3.22 Y-1.22 Z0.8 E0.14018
G1 X -3.22 Y-1.71 Z0.8 E0.14768
G1 X -3.22 Y-2.2 Z0.8 E0.15518
G1 X -3.22 Y-2.69 Z0.8 E0.16267
G1 X -3.22 Y-3.18 Z0.8 E0.17017
G1 X -3.22 Y-3.67 Z0.8 E0.17767
G1 X -3.22 Y-4.15 Z0.8 E0.18517
G1 X -3.22 Y-4.64 Z0.8 E0.19267
G1 X-3.219 Y-5.132 E0.20017 ; infill
G1 X -3.56 Y-4.77 Z0.8 E0.17647
G1 X -3.9 Y-4.41 Z0.8 E0.15277
G1 X -4.23 Y-4.04 Z0.8 E0.12907
G1 X -4.57 Y-3.68 Z0.8 E0.10538
G1 X -4.91 Y-3.32 Z0.8 E0.08168
G1 X-5.247 Y-2.957 E0.05798 ; infill
```

```
G1 X -5.25 Y-2.46 Z0.8 E0.06276
G1 X -5.25 Y-1.97 Z0.8 E0.06754
G1 X -5.25 Y-1.48 Z0.8 E0.07232
G1 X -5.25 Y-0.99 Z0.8 E0.0771
G1 X -5.25 Y-0.49 Z0.8 E0.08188
G1 X -5.25 Y0.0 Z0.8 E0.08666
G1 X -5.25 Y0.49 Z0.8 E0.09145
G1 X -5.25 Y0.99 Z0.8 E0.09623
G1 X -5.25 Y1.48 Z0.8 E0.10101
G1 X -5.25 Y1.97 Z0.8 E0.10579
G1 X -5.25 Y2.46 Z0.8 E0.11057
G1 X-5.247 Y2.957 E0.11535 ; infill
M761 ; Deactivate valve for PH1
G1 E-2.00000 F300.00000 ; retract
G0 Z30
M84 ;disable motors
;END
```

**<u>Model 1(rectangular structure)</u>**

```
; external perimeters extrusion width = 0.45mm
; perimeters extrusion width = 0.48mm
; infill extrusion width = 0.48mm
; solid infill extrusion width = 0.48mm
; top infill extrusion width = 0.48mm
; support material extrusion width = 0.45mm
G21 ; set units to millimeters
G90 ; use absolute coordinates
M83 ; use relative distances for extrusion
G1 Z0.400 F600.000 ; move to next layer (0)
G1 E-2.00000 F2400.00000 ; retract
G1 X-2.274 Y1.307 F600.000 ; move to first perimeter point
M760
G4 P0
G1 E2.00000 F2400.00000 ; unretract
G1 X-2.467 Y1.277 E0.00347 F600.000 ; perimeter
G1 X-2.659 Y1.182 E0.00379 ; perimeter
G1 X-2.825 Y1.038 E0.00391 ; perimeter
G1 X-2.963 Y0.854 E0.00409 ; perimeter
G1 X-3.071 Y0.639 E0.00429 ; perimeter
G1 X-3.149 Y0.399 E0.00447 ; perimeter
G1 X-3.197 Y0.140 E0.00468 ; perimeter
G1 X-3.209 Y-0.249 E0.00693 ; perimeter
G1 X-3.177 Y-0.513 E0.00472 ; perimeter
G1 X-3.114 Y-0.765 E0.00463 ; perimeter
G1 X-2.965 Y-1.090 E0.00635 ; perimeter
G1 X-2.825 Y-1.278 E0.00416 ; perimeter
G1 X-2.659 Y-1.422 E0.00391 ; perimeter
G1 X-2.467 Y-1.517 E0.00379 ; perimeter
G1 X-2.275 Y-1.547 E0.00347 ; perimeter
G1 X-1.822 Y-1.547 E1
G1 X-1.369 Y-1.547 E1
```

```
G1 X-0.916 Y-1.547 E1
G1 X-0.463 Y-1.547 E1
G1 X-0.01 Y-1.547 E1
G1 X0.443 Y-1.547 E1
G1 X0.896 Y-1.547 E1
G1 X1.349 Y-1.547 E1
G1 X1.802 Y-1.547 E1
G1 X2.255 Y-1.547 E0.08052 ; perimeter
G1 X2.447 Y-1.517 E0.00347 ; perimeter
G1 X2.639 Y-1.422 E0.00379 ; perimeter
G1 X2.805 Y-1.278 E0.00391 ; perimeter
G1 X2.945 Y-1.090 E0.00416 ; perimeter
G1 X3.095 Y-0.763 E0.00641 ; perimeter
G1 X3.177 Y-0.386 E0.00685 ; perimeter
G1 X3.192 Y-0.120 E0.00474 ; perimeter
G1 X3.177 Y0.146 E0.00474 ; perimeter
G1 X3.095 Y0.523 E0.00685 ; perimeter
G1 X2.945 Y0.850 E0.00641 ; perimeter
G1 X2.802 Y1.043 E0.00426 ; perimeter
G1 X2.553 Y1.235 E0.00559 ; perimeter
G1 X2.345 Y1.302 E0.00388 ; perimeter
G1 X1.8898 Y1.3025 E1
G1 X1.4346 Y1.303 E1
G1 X0.9794 Y1.3035 E1
G1 X0.5242 Y1.304 E1
G1 X0.069 Y1.3045 E1
G1 X-0.3862 Y1.305 E1
G1 X-0.8414 Y1.3055 E1
G1 X-1.2966 Y1.306 E1
G1 X-1.7518 Y1.3065 E1
G1 X-2.207 Y1.307 E0.08092 ; perimeter
M761
G1 X-3.135 Y-1.892 F600.000 ; move to first perimeter point
```

```
M760
G4 P0
G1 X-2.671 Y-0.2925 E1
G1 E2.00000 F2400.00000 ; unretract
G1 X-3.005 Y-1.920 E0.00237 F600.000 ; perimeter
G1 X-2.50658333333 Y-1.91991666667 E1
G1 X-2.00816666667 Y-1.91983333333 E1
G1 X-1.50975 Y-1.91975 E1
G1 X-1.01133333333 Y-1.91966666667 E1
G1 X-0.512916666667 Y-1.91958333333 E1
G1 X-0.0145 Y-1.9195 E1
G1 X0.483916666667 Y-1.91941666667 E1
G1 X0.982333333333 Y-1.91933333333 E1
G1 X1.48075 Y-1.91925 E1
G1 X1.97916666667 Y-1.91916666667 E1
G1 X2.47758333333 Y-1.91908333333 E1
G1 X2.976 Y-1.919 E0.10633 ; perimeter
G1 X3.108 Y-1.893 E0.00240 ; perimeter
G1 X3.243 Y-1.811 E0.00280 ; perimeter
G1 X3.384 Y-1.661 E0.00366 ; perimeter
G1 X3.517 Y-1.443 E0.00454 ; perimeter
G1 X3.631 Y-1.167 E0.00532 ; perimeter
G1 X3.717 Y-0.846 E0.00589 ; perimeter
G1 X3.751 Y-0.576 E1
G1 X3.785 Y-0.306 E0.00968 ; perimeter
G1 X3.778 Y-0.0265 E1
G1 X3.771 Y0.253 E0.00993 ; perimeter
G1 X3.718 Y0.604 E0.00632 ; perimeter
G1 X3.631 Y0.927 E0.00594 ; perimeter
G1 X3.517 Y1.203 E0.00532 ; perimeter
G1 X3.384 Y1.421 E0.00454 ; perimeter
G1 X3.238 Y1.577 E0.00380 ; perimeter
G1 X3.116 Y1.651 E0.00253 ; perimeter
```

```
G1 X2.976 Y1.680 E0.00255 ; perimeter
G1 X2.47758333333 Y1.68 E1
G1 X1.97916666667 Y1.68 E1
G1 X1.48075 Y1.68 E1
G1 X0.982333333333 Y1.68 E1
G1 X0.483916666667 Y1.68 E1
G1 X-0.0145 Y1.68 E1
G1 X-0.512916666667 Y1.68 E1
G1 X-1.01133333333 Y1.68 E1
G1 X-1.50975 Y1.68 E1
G1 X-2.00816666667 Y1.68 E1
G1 X-2.50658333333 Y1.68 E1
G1 X-3.005 Y1.680 E0.10632 ; perimeter
G1 X-3.127 Y1.654 E0.00222 ; perimeter
G1 X-3.263 Y1.571 E0.00283 ; perimeter
G1 X-3.404 Y1.421 E0.00366 ; perimeter
G1 X-3.537 Y1.203 E0.00454 ; perimeter
G1 X-3.650 Y0.929 E0.00527 ; perimeter
G1 X-3.7095 Y0.6795 E1
G1 X-3.769 Y0.430 E0.00912 ; perimeter
G1 X-3.7895 Y0.1535 E1
G1 X-3.810 Y-0.123 E0.00985 ; perimeter
G1 X-3.792 Y-0.490 E0.00654 ; perimeter
G1 X-3.738 Y-0.841 E0.00631 ; perimeter
G1 X-3.597 Y-1.311 E0.00873 ; perimeter
G1 X-3.401 Y-1.664 E0.00718 ; perimeter
G1 X-3.195 Y-1.861 E0.00507 ; perimeter
M761
G1 X-2.9085 Y-1.812 E1
G1 X-2.622 Y-1.763 F600.000 ; move to first infill point
M760
G4 P0
G1 X-3.030 Y-1.557 E0.01846 F600.000 ; infill
```

```
G1 X-3.120 Y-1.459 E0.00537 ; infill
G1 X-3.258 Y-1.243 E0.01038 ; infill
G1 X-3.3455 Y-1.0075 E1
G1 X-3.433 Y-0.772 E0.02029 ; infill
G1 X-3.489 Y-0.460 E0.01282 ; infill
G1 X-3.507 Y-0.126 E0.01350 ; infill
G1 X-3.490 Y0.183 E0.01251 ; infill
G1 X-3.447 Y0.465 E0.01155 ; infill
G1 X-3.361 Y0.771 E0.01284 ; infill
G1 X-3.240 Y1.043 E0.01203 ; infill
G1 X-3.045 Y1.302 E0.01309 ; infill
G1 X-2.622 Y1.523 E0.01929 ; infill
M761
G1 X-2.591 Y1.461 F600.000 ; move to first infill point
M760
G4 P0
G1 X-2.287 Y1.494 E0.00544 F600.000 ; infill
G1 X-1.8213 Y1.4937 E1
G1 X-1.3556 Y1.4934 E1
G1 X-0.8899 Y1.4931 E1
G1 X-0.4242 Y1.4928 E1
G1 X0.0415 Y1.4925 E1
G1 X0.5072 Y1.4922 E1
G1 X0.9729 Y1.4919 E1
G1 X1.4386 Y1.4916 E1
G1 X1.9043 Y1.4913 E1
G1 X2.370 Y1.491 E0.08278 ; infill
G1 X2.672 Y1.444 E0.00545 ; infill
M761
G1 X2.684 Y1.469 F600.000 ; move to first infill point
M760
G4 P0
G1 X3.010 Y1.317 E0.01454 F600.000 ; infill
```

```
G1 X3.108 Y1.210 E0.00584 ; infill
G1 X3.240 Y1.004 E0.00992 ; infill
G1 X3.411 Y0.540 E0.01999 ; infill
G1 X3.473 Y0.198 E0.01402 ; infill
G1 X3.484 Y-0.291 E0.01976 ; infill
G1 X3.4395 Y-0.568 E1
G1 X3.395 Y-0.845 E0.02269 ; infill
G1 X3.216 Y-1.290 E0.01940 ; infill
G1 X3.025 Y-1.542 E0.01276 ; infill
G1 X2.594 Y-1.762 E0.01955 ; infill
M761
G1 X2.569 Y-1.710 F600.000 ; move to first infill point
M760
G4 P0
G1 X2.267 Y-1.733 E0.00538 F600.000 ; infill
G1 X1.8116 Y-1.733 E1
G1 X1.3562 Y-1.733 E1
G1 X0.9008 Y-1.733 E1
G1 X0.4454 Y-1.733 E1
G1 X-0.01 Y-1.733 E1
G1 X-0.4654 Y-1.733 E1
G1 X-0.9208 Y-1.733 E1
G1 X-1.3762 Y-1.733 E1
G1 X-1.8316 Y-1.733 E1
G1 X-2.287 Y-1.733 E0.08095 ; infill
G1 X-2.591 Y-1.700 E0.00544 ; infill
M761
G1 Z0.800 F600.000 ; move to next layer (1)
G1 X-2.672 Y-1.443 F600.000 ; move to first perimeter point
M760
G4 P0
G1 X-2.5735 Y-1.492 E1
G1 X-2.475 Y-1.541 E0.00454 F240.000 ; perimeter
```

```
G1 X-2.3755 Y-1.5565 E1
G1 X-2.276 Y-1.572 E0.00414 ; perimeter
G1 X-2.07895652174 Y-1.572 E1
G1 X-1.88191304348 Y-1.572 E1
G1 X-1.68486956522 Y-1.572 E1
G1 X-1.48782608696 Y-1.572 E1
G1 X-1.2907826087 Y-1.572 E1
G1 X-1.09373913043 Y-1.572 E1
G1 X-0.896695652174 Y-1.572 E1
G1 X-0.699652173913 Y-1.572 E1
G1 X-0.502608695652 Y-1.572 E1
G1 X-0.305565217391 Y-1.572 E1
G1 X-0.10852173913 Y-1.572 E1
G1 X0.0885217391304 Y-1.572 E1
G1 X0.285565217391 Y-1.572 E1
G1 X0.482608695652 Y-1.572 E1
G1 X0.679652173913 Y-1.572 E1
G1 X0.876695652174 Y-1.572 E1
G1 X1.07373913043 Y-1.572 E1
G1 X1.2707826087 Y-1.572 E1
G1 X1.46782608696 Y-1.572 E1
G1 X1.66486956522 Y-1.572 E1
G1 X1.86191304348 Y-1.572 E1
G1 X2.05895652174 Y-1.572 E1
G1 X2.256 Y-1.572 E0.09341 ; perimeter
G1 X2.3555 Y-1.5565 E1
G1 X2.455 Y-1.541 E0.00414 ; perimeter
G1 X2.5535 Y-1.492 E1
G1 X2.652 Y-1.443 E0.00454 ; perimeter
G1 X2.738 Y-1.369 E1
G1 X2.824 Y-1.295 E0.00466 ; perimeter
G1 X2.8955 Y-1.199 E1
G1 X2.967 Y-1.103 E0.00493 ; perimeter
```

```
G1 X3.043 Y-0.9365 E1
G1 X3.119 Y-0.770 E0.00754 ; perimeter
G1 X3.1605 Y-0.58 E1
G1 X3.202 Y-0.390 E0.00803 ; perimeter
G1 X3.2095 Y-0.255 E1
G1 X3.217 Y-0.120 E0.00557 ; perimeter
G1 X3.2095 Y0.015 E1
G1 X3.202 Y0.150 E0.00557 ; perimeter
G1 X3.1605 Y0.34 E1
G1 X3.119 Y0.530 E0.00803 ; perimeter
G1 X3.043 Y0.6965 E1
G1 X2.967 Y0.863 E0.00754 ; perimeter
G1 X2.8935 Y0.9615 E1
G1 X2.820 Y1.060 E0.00506 ; perimeter
G1 X2.692 Y1.1585 E1
G1 X2.564 Y1.257 E0.00664 ; perimeter
G1 X2.4565 Y1.292 E1
G1 X2.349 Y1.327 E0.00467 ; perimeter
G1 X2.15629166667 Y1.32720833333 E1
G1 X1.96358333333 Y1.32741666667 E1
G1 X1.770875 Y1.327625 E1
G1 X1.57816666667 Y1.32783333333 E1
G1 X1.38545833333 Y1.32804166667 E1
G1 X1.19275 Y1.32825 E1
G1 X1.00004166667 Y1.32845833333 E1
G1 X0.807333333333 Y1.32866666667 E1
G1 X0.614625 Y1.328875 E1
G1 X0.421916666667 Y1.32908333333 E1
G1 X0.229208333333 Y1.32929166667 E1
G1 X0.0365 Y1.3295 E1
G1 X-0.156208333333 Y1.32970833333 E1
G1 X-0.348916666667 Y1.32991666667 E1
G1 X-0.541625 Y1.330125 E1
```

```
G1 X-0.734333333333 Y1.33033333333 E1
G1 X-0.927041666667 Y1.33054166667 E1
G1 X-1.11975 Y1.33075 E1
G1 X-1.31245833333 Y1.33095833333 E1
G1 X-1.50516666667 Y1.33116666667 E1
G1 X-1.697875 Y1.331375 E1
G1 X-1.89058333333 Y1.33158333333 E1
G1 X-2.08329166667 Y1.33179166667 E1
G1 X-2.276 Y1.332 E0.09531 ; perimeter
G1 X-2.3755 Y1.3165 E1
G1 X-2.475 Y1.301 E0.00414 ; perimeter
G1 X-2.5735 Y1.252 E1
G1 X-2.672 Y1.203 E0.00454 ; perimeter
G1 X-2.758 Y1.129 E1
G1 X-2.844 Y1.055 E0.00466 ; perimeter
G1 X-2.914 Y0.961 E1
G1 X-2.984 Y0.867 E0.00484 ; perimeter
G1 X-3.0395 Y0.7575 E1
G1 X-3.095 Y0.648 E0.00505 ; perimeter
G1 X-3.1345 Y0.527 E1
G1 X-3.174 Y0.406 E0.00525 ; perimeter
G1 X-3.198 Y0.2745 E1
G1 X-3.222 Y0.143 E0.00550 ; perimeter
G1 X-3.228 Y-0.0535 E1
G1 X-3.234 Y-0.250 E0.00811 ; perimeter
G1 X-3.218 Y-0.3835 E1
G1 X-3.202 Y-0.517 E0.00554 ; perimeter
G1 X-3.1695 Y-0.6455 E1
G1 X-3.137 Y-0.774 E0.00545 ; perimeter
G1 X-3.062 Y-0.9385 E1
G1 X-2.987 Y-1.103 E0.00746 ; perimeter
G1 X-2.9155 Y-1.199 E1
G1 X-2.844 Y-1.295 E0.00493 ; perimeter
```

```
G1 X-2.724 Y-1.399 E0.00327 ; perimeter
M761
G1 X-2.9255 Y-1.6335 E1
G1 X-3.127 Y-1.868 F600.000 ; move to first perimeter point
M760
G4 P0
G1 X-3.002 Y-1.895 E0.00263 F240.000 ; perimeter
G1 X-2.8028 Y-1.89496666667 E1
G1 X-2.6036 Y-1.89493333333 E1
G1 X-2.4044 Y-1.8949 E1
G1 X-2.2052 Y-1.89486666667 E1
G1 X-2.006 Y-1.89483333333 E1
G1 X-1.8068 Y-1.8948 E1
G1 X-1.6076 Y-1.89476666667 E1
G1 X-1.4084 Y-1.89473333333 E1
G1 X-1.2092 Y-1.8947 E1
G1 X-1.01 Y-1.89466666667 E1
G1 X-0.8108 Y-1.89463333333 E1
G1 X-0.6116 Y-1.8946 E1
G1 X-0.4124 Y-1.89456666667 E1
G1 X-0.2132 Y-1.89453333333 E1
G1 X-0.014 Y-1.8945 E1
G1 X0.1852 Y-1.89446666667 E1
G1 X0.3844 Y-1.89443333333 E1
G1 X0.5836 Y-1.8944 E1
G1 X0.7828 Y-1.89436666667 E1
G1 X0.982 Y-1.89433333333 E1
G1 X1.1812 Y-1.8943 E1
G1 X1.3804 Y-1.89426666667 E1
G1 X1.5796 Y-1.89423333333 E1
G1 X1.7788 Y-1.8942 E1
G1 X1.978 Y-1.89416666667 E1
G1 X2.1772 Y-1.89413333333 E1
```

```
G1 X2.3764 Y-1.8941 E1
G1 X2.5756 Y-1.89406666667 E1
G1 X2.7748 Y-1.89403333333 E1
G1 X2.974 Y-1.894 E0.12315 ; perimeter
G1 X3.099 Y-1.869 E0.00263 ; perimeter
G1 X3.227 Y-1.792 E0.00308 ; perimeter
G1 X3.2955 Y-1.719 E1
G1 X3.364 Y-1.646 E0.00412 ; perimeter
G1 X3.4295 Y-1.539 E1
G1 X3.495 Y-1.432 E0.00517 ; perimeter
G1 X3.551 Y-1.2955 E1
G1 X3.607 Y-1.159 E0.00609 ; perimeter
G1 X3.65 Y-1.0005 E1
G1 X3.693 Y-0.842 E0.00676 ; perimeter
G1 X3.71533333333 Y-0.663 E1
G1 X3.73766666667 Y-0.484 E1
G1 X3.760 Y-0.305 E0.01115 ; perimeter
G1 X3.75533333333 Y-0.12 E1
G1 X3.75066666667 Y0.065 E1
G1 X3.746 Y0.250 E0.01144 ; perimeter
G1 X3.7195 Y0.4245 E1
G1 X3.693 Y0.599 E0.00726 ; perimeter
G1 X3.65 Y0.759 E1
G1 X3.607 Y0.919 E0.00683 ; perimeter
G1 X3.551 Y1.0555 E1
G1 X3.495 Y1.192 E0.00609 ; perimeter
G1 X3.4295 Y1.299 E1
G1 X3.364 Y1.406 E0.00517 ; perimeter
G1 X3.2925 Y1.482 E1
G1 X3.221 Y1.558 E0.00430 ; perimeter
G1 X3.108 Y1.627 E0.00274 ; perimeter
G1 X2.973 Y1.655 E0.00283 ; perimeter
G1 X2.77383333333 Y1.655 E1
```

```
G1 X2.57466666667 Y1.655 E1
G1 X2.3755 Y1.655 E1
G1 X2.17633333333 Y1.655 E1
G1 X1.97716666667 Y1.655 E1
G1 X1.778 Y1.655 E1
G1 X1.57883333333 Y1.655 E1
G1 X1.37966666667 Y1.655 E1
G1 X1.1805 Y1.655 E1
G1 X0.981333333333 Y1.655 E1
G1 X0.782166666667 Y1.655 E1
G1 X0.583 Y1.655 E1
G1 X0.383833333333 Y1.655 E1
G1 X0.184666666667 Y1.655 E1
G1 X-0.0145 Y1.655 E1
G1 X-0.213666666667 Y1.655 E1
G1 X-0.412833333333 Y1.655 E1
G1 X-0.612 Y1.655 E1
G1 X-0.811166666667 Y1.655 E1
G1 X-1.01033333333 Y1.655 E1
G1 X-1.2095 Y1.655 E1
G1 X-1.40866666667 Y1.655 E1
G1 X-1.60783333333 Y1.655 E1
G1 X-1.807 Y1.655 E1
G1 X-2.00616666667 Y1.655 E1
G1 X-2.20533333333 Y1.655 E1
G1 X-2.4045 Y1.655 E1
G1 X-2.60366666667 Y1.655 E1
G1 X-2.80283333333 Y1.655 E1
G1 X-3.002 Y1.655 E0.12313 ; perimeter
G1 X-3.118 Y1.630 E0.00244 ; perimeter
G1 X-3.247 Y1.552 E0.00312 ; perimeter
G1 X-3.3155 Y1.479 E1
G1 X-3.384 Y1.406 E0.00412 ; perimeter
```

```
G1 X-3.4495 Y1.299 E1
G1 X-3.515 Y1.192 E0.00517 ; perimeter
G1 X-3.5705 Y1.057 E1
G1 X-3.626 Y0.922 E0.00602 ; perimeter
G1 X-3.66533333333 Y0.756666666667 E1
G1 X-3.70466666667 Y0.591333333333 E1
G1 X-3.744 Y0.426 E0.01049 ; perimeter
G1 X-3.75766666667 Y0.243 E1
G1 X-3.77133333333 Y0.06 E1
G1 X-3.785 Y-0.123 E0.01135 ; perimeter
G1 X-3.776 Y-0.3055 E1
G1 X-3.767 Y-0.488 E0.00752 ; perimeter
G1 X-3.7405 Y-0.6615 E1
G1 X-3.714 Y-0.835 E0.00725 ; perimeter
G1 X-3.66733333333 Y-0.990333333333 E1
G1 X-3.62066666667 Y-1.14566666667 E1
G1 X-3.574 Y-1.301 E0.01002 ; perimeter
G1 X-3.4775 Y-1.475 E1
G1 X-3.381 Y-1.649 E0.00820 ; perimeter
G1 X-3.284 Y-1.743 E1
G1 X-3.187 Y-1.837 E0.00556 ; perimeter
M761
G1 X-2.9045 Y-1.765 E1
G1 X-2.622 Y-1.693 F600.000 ; move to first infill point
M760
G4 P0
G1 X-2.816 Y-1.647 E0.00451 F240.000 ; infill
G1 X-2.917 Y-1.606 E1
G1 X-3.018 Y-1.565 E0.00490 ; infill
G1 X-3.120 Y-1.459 E0.00331 ; infill
G1 X-3.1875 Y-1.3535 E1
G1 X-3.255 Y-1.248 E0.00564 ; infill
G1 X-3.31433333333 Y-1.08933333333 E1
```

G1 X-3.37366666667 Y-0.930666666667 E1

G1 X-3.433 Y-0.772 E0.01144 ; infill

G1 X-3.461 Y-0.616 E1

G1 X-3.489 Y-0.460 E0.00714 ; infill

G1 X-3.498 Y-0.293 E1

G1 X-3.507 Y-0.126 E0.00751 ; infill

G1 X-3.499 Y0.026 E1

G1 X-3.491 Y0.178 E0.00686 ; infill

G1 X-3.4685 Y0.3235 E1

G1 X-3.446 Y0.469 E0.00662 ; infill

G1 X-3.404 Y0.618 E1

G1 X-3.362 Y0.767 E0.00696 ; infill

G1 X-3.302 Y0.903 E1

G1 X-3.242 Y1.039 E0.00669 ; infill

G1 X-3.185 Y1.1245 E1

G1 X-3.128 Y1.210 E0.00464 ; infill

G1 X-3.000 Y1.335 E0.00401 ; infill

G1 X-2.817 Y1.407 E0.00444 ; infill

G1 X-2.719 Y1.43 E1

G1 X-2.621 Y1.453 E0.00451 ; infill

M761

G1 E-2.00000 F2400.00000 ; retract

G1 X-2.14463636364 Y1.45290909091 E1

G1 X-1.66827272727 Y1.45281818182 E1

G1 X-1.19190909091 Y1.45272727273 E1

G1 X-0.715545454545 Y1.45263636364 E1

G1 X-0.239181818182 Y1.45254545455 E1

G1 X0.237181818182 Y1.45245454545 E1

G1 X0.713545454545 Y1.45236363636 E1

G1 X1.18990909091 Y1.45227272727 E1

G1 X1.66627272727 Y1.45218181818 E1

G1 X2.14263636364 Y1.45209090909 E1

G1 X2.619 Y1.452 F600.000 ; move to first infill point

```
M760
G4 P0
G1 X-0.874333333333 Y1.45266666667 E1
G1 X0.872333333333 Y1.45233333333 E1
G1 E2.00000 F2400.00000 ; unretract
G1 X2.746 Y1.40933333333 E1
G1 X2.873 Y1.36666666667 E1
G1 X3.000 Y1.324 E0.00904 F240.000 ; infill
G1 X3.108 Y1.210 E0.00352 ; infill
G1 X3.1675 Y1.121 E1
G1 X3.227 Y1.032 E0.00483 ; infill
G1 X3.2735 Y0.9265 E1
G1 X3.320 Y0.821 E0.00518 ; infill
G1 X3.3655 Y0.6805 E1
G1 X3.411 Y0.540 E0.00666 ; infill
G1 X3.442 Y0.367 E1
G1 X3.473 Y0.194 E0.00791 ; infill
G1 X3.4795 Y0.0345 E1
G1 X3.486 Y-0.125 E0.00716 ; infill
G1 X3.479 Y-0.2725 E1
G1 X3.472 Y-0.420 E0.00666 ; infill
G1 X3.441 Y-0.603 E1
G1 X3.410 Y-0.786 E0.00835 ; infill
G1 X3.365 Y-0.9235 E1
G1 X3.320 Y-1.061 E0.00651 ; infill
G1 X3.2735 Y-1.1665 E1
G1 X3.227 Y-1.272 E0.00518 ; infill
G1 X3.1675 Y-1.361 E1
G1 X3.108 Y-1.450 E0.00483 ; infill
G1 X2.984 Y-1.572 E0.00391 ; infill
G1 X2.89 Y-1.609 E1
G1 X2.796 Y-1.646 E0.00454 ; infill
G1 X2.603 Y-1.692 E0.00446 ; infill
```

```
M761
G1 Z1.200 F600.000 ; move to next layer (2)
G1 E-2.00000 F2400.00000 ; retract
G1 X2.12345454545 Y-1.66936363636 E1
G1 X1.64390909091 Y-1.64672727273 E1
G1 X1.16436363636 Y-1.62409090909 E1
G1 X0.684818181818 Y-1.60145454545 E1
G1 X0.205272727273 Y-1.57881818182 E1
G1 X-0.274272727273 Y-1.55618181818 E1
G1 X-0.753818181818 Y-1.53354545455 E1
G1 X-1.23336363636 Y-1.51090909091 E1
G1 X-1.71290909091 Y-1.48827272727 E1
G1 X-2.19245454545 Y-1.46563636364 E1
G1 X-2.672 Y-1.443 F600.000 ; move to first perimeter point
M760
G4 P0
G1 X0.844666666667 Y-1.609 E1
G1 X-0.913666666667 Y-1.526 E1
G1 E2.00000 F2400.00000 ; unretract
G1 X-2.5735 Y-1.492 E1
G1 X-2.475 Y-1.541 E0.00454 F240.000 ; perimeter
G1 X-2.3755 Y-1.5565 E1
G1 X-2.276 Y-1.572 E0.00414 ; perimeter
G1 X-2.07895652174 Y-1.572 E1
G1 X-1.88191304348 Y-1.572 E1
G1 X-1.68486956522 Y-1.572 E1
G1 X-1.48782608696 Y-1.572 E1
G1 X-1.2907826087 Y-1.572 E1
G1 X-1.09373913043 Y-1.572 E1
G1 X-0.896695652174 Y-1.572 E1
G1 X-0.699652173913 Y-1.572 E1
G1 X-0.502608695652 Y-1.572 E1
G1 X-0.305565217391 Y-1.572 E1
```

```
G1 X-0.10852173913 Y-1.572 E1
G1 X0.0885217391304 Y-1.572 E1
G1 X0.285565217391 Y-1.572 E1
G1 X0.482608695652 Y-1.572 E1
G1 X0.679652173913 Y-1.572 E1
G1 X0.876695652174 Y-1.572 E1
G1 X1.07373913043 Y-1.572 E1
G1 X1.2707826087 Y-1.572 E1
G1 X1.46782608696 Y-1.572 E1
G1 X1.66486956522 Y-1.572 E1
G1 X1.86191304348 Y-1.572 E1
G1 X2.05895652174 Y-1.572 E1
G1 X2.256 Y-1.572 E0.09341 ; perimeter
G1 X2.3555 Y-1.5565 E1
G1 X2.455 Y-1.541 E0.00414 ; perimeter
G1 X2.5535 Y-1.492 E1
G1 X2.652 Y-1.443 E0.00454 ; perimeter
G1 X2.738 Y-1.369 E1
G1 X2.824 Y-1.295 E0.00466 ; perimeter
G1 X2.8955 Y-1.199 E1
G1 X2.967 Y-1.103 E0.00493 ; perimeter
G1 X3.043 Y-0.9365 E1
G1 X3.119 Y-0.770 E0.00754 ; perimeter
G1 X3.1605 Y-0.58 E1
G1 X3.202 Y-0.390 E0.00803 ; perimeter
G1 X3.2095 Y-0.255 E1
G1 X3.217 Y-0.120 E0.00557 ; perimeter
G1 X3.2095 Y0.015 E1
G1 X3.202 Y0.150 E0.00557 ; perimeter
G1 X3.1605 Y0.34 E1
G1 X3.119 Y0.530 E0.00803 ; perimeter
G1 X3.043 Y0.6965 E1
G1 X2.967 Y0.863 E0.00754 ; perimeter
```

G1 X2.8935 Y0.9615 E1

G1 X2.820 Y1.060 E0.00506 ; perimeter

G1 X2.692 Y1.1585 E1

G1 X2.564 Y1.257 E0.00664 ; perimeter

G1 X2.4565 Y1.292 E1

G1 X2.349 Y1.327 E0.00467 ; perimeter

G1 X2.15629166667 Y1.32720833333 E1

G1 X1.96358333333 Y1.32741666667 E1

G1 X1.770875 Y1.327625 E1

G1 X1.57816666667 Y1.32783333333 E1

G1 X1.38545833333 Y1.32804166667 E1

G1 X1.19275 Y1.32825 E1

G1 X1.00004166667 Y1.32845833333 E1

G1 X0.807333333333 Y1.32866666667 E1

G1 X0.614625 Y1.328875 E1

G1 X0.421916666667 Y1.32908333333 E1

G1 X0.229208333333 Y1.32929166667 E1

G1 X0.0365 Y1.3295 E1

G1 X-0.156208333333 Y1.32970833333 E1

G1 X-0.348916666667 Y1.32991666667 E1

G1 X-0.541625 Y1.330125 E1

G1 X-0.734333333333 Y1.33033333333 E1

G1 X-0.927041666667 Y1.33054166667 E1

G1 X-1.11975 Y1.33075 E1

G1 X-1.31245833333 Y1.33095833333 E1

G1 X-1.50516666667 Y1.33116666667 E1

G1 X-1.697875 Y1.331375 E1

G1 X-1.89058333333 Y1.33158333333 E1

G1 X-2.08329166667 Y1.33179166667 E1

G1 X-2.276 Y1.332 E0.09531 ; perimeter

G1 X-2.3755 Y1.3165 E1

G1 X-2.475 Y1.301 E0.00414 ; perimeter

G1 X-2.5735 Y1.252 E1

G1 X-2.672 Y1.203 E0.00454 ; perimeter

G1 X-2.758 Y1.129 E1

G1 X-2.844 Y1.055 E0.00466 ; perimeter

G1 X-2.914 Y0.961 E1

G1 X-2.984 Y0.867 E0.00484 ; perimeter

G1 X-3.0395 Y0.7575 E1

G1 X-3.095 Y0.648 E0.00505 ; perimeter

G1 X-3.1345 Y0.527 E1

G1 X-3.174 Y0.406 E0.00525 ; perimeter

G1 X-3.198 Y0.2745 E1

G1 X-3.222 Y0.143 E0.00550 ; perimeter

G1 X-3.228 Y-0.0535 E1

G1 X-3.234 Y-0.250 E0.00811 ; perimeter

G1 X-3.218 Y-0.3835 E1

G1 X-3.202 Y-0.517 E0.00554 ; perimeter

G1 X-3.1695 Y-0.6455 E1

G1 X-3.137 Y-0.774 E0.00545 ; perimeter

G1 X-3.062 Y-0.9385 E1

G1 X-2.987 Y-1.103 E0.00746 ; perimeter

G1 X-2.9155 Y-1.199 E1

G1 X-2.844 Y-1.295 E0.00493 ; perimeter

G1 X-2.724 Y-1.399 E0.00327 ; perimeter

M761

G1 X-2.9255 Y-1.6335 E1

G1 X-3.127 Y-1.868 F600.000 ; move to first perimeter point

M760

G4 P0

G1 X-3.002 Y-1.895 E0.00263 F240.000 ; perimeter

G1 X-2.8028 Y-1.89496666667 E1

G1 X-2.6036 Y-1.89493333333 E1

G1 X-2.4044 Y-1.8949 E1

G1 X-2.2052 Y-1.89486666667 E1

G1 X-2.006 Y-1.89483333333 E1

```
G1 X-1.8068 Y-1.8948 E1
G1 X-1.6076 Y-1.89476666667 E1
G1 X-1.4084 Y-1.89473333333 E1
G1 X-1.2092 Y-1.8947 E1
G1 X-1.01 Y-1.89466666667 E1
G1 X-0.8108 Y-1.89463333333 E1
G1 X-0.6116 Y-1.8946 E1
G1 X-0.4124 Y-1.89456666667 E1
G1 X-0.2132 Y-1.89453333333 E1
G1 X-0.014 Y-1.8945 E1
G1 X0.1852 Y-1.89446666667 E1
G1 X0.3844 Y-1.89443333333 E1
G1 X0.5836 Y-1.8944 E1
G1 X0.7828 Y-1.89436666667 E1
G1 X0.982 Y-1.89433333333 E1
G1 X1.1812 Y-1.8943 E1
G1 X1.3804 Y-1.89426666667 E1
G1 X1.5796 Y-1.89423333333 E1
G1 X1.7788 Y-1.8942 E1
G1 X1.978 Y-1.89416666667 E1
G1 X2.1772 Y-1.89413333333 E1
G1 X2.3764 Y-1.8941 E1
G1 X2.5756 Y-1.89406666667 E1
G1 X2.7748 Y-1.89403333333 E1
G1 X2.974 Y-1.894 E0.12315 ; perimeter
G1 X3.099 Y-1.869 E0.00263 ; perimeter
G1 X3.227 Y-1.792 E0.00308 ; perimeter
G1 X3.2955 Y-1.719 E1
G1 X3.364 Y-1.646 E0.00412 ; perimeter
G1 X3.4295 Y-1.539 E1
G1 X3.495 Y-1.432 E0.00517 ; perimeter
G1 X3.551 Y-1.2955 E1
G1 X3.607 Y-1.159 E0.00609 ; perimeter
```

G1 X3.65 Y-1.0005 E1

G1 X3.693 Y-0.842 E0.00676 ; perimeter

G1 X3.71533333333 Y-0.663 E1

G1 X3.73766666667 Y-0.484 E1

G1 X3.760 Y-0.305 E0.01115 ; perimeter

G1 X3.75533333333 Y-0.12 E1

G1 X3.75066666667 Y0.065 E1

G1 X3.746 Y0.250 E0.01144 ; perimeter

G1 X3.7195 Y0.4245 E1

G1 X3.693 Y0.599 E0.00726 ; perimeter

G1 X3.65 Y0.759 E1

G1 X3.607 Y0.919 E0.00683 ; perimeter

G1 X3.551 Y1.0555 E1

G1 X3.495 Y1.192 E0.00609 ; perimeter

G1 X3.4295 Y1.299 E1

G1 X3.364 Y1.406 E0.00517 ; perimeter

G1 X3.2925 Y1.482 E1

G1 X3.221 Y1.558 E0.00430 ; perimeter

G1 X3.108 Y1.627 E0.00274 ; perimeter

G1 X2.973 Y1.655 E0.00283 ; perimeter

G1 X2.77383333333 Y1.655 E1

G1 X2.57466666667 Y1.655 E1

G1 X2.3755 Y1.655 E1

G1 X2.17633333333 Y1.655 E1

G1 X1.97716666667 Y1.655 E1

G1 X1.778 Y1.655 E1

G1 X1.57883333333 Y1.655 E1

G1 X1.37966666667 Y1.655 E1

G1 X1.1805 Y1.655 E1

G1 X0.981333333333 Y1.655 E1

G1 X0.782166666667 Y1.655 E1

G1 X0.583 Y1.655 E1

G1 X0.383833333333 Y1.655 E1

G1 X0.184666666667 Y1.655 E1

G1 X-0.0145 Y1.655 E1

G1 X-0.213666666667 Y1.655 E1

G1 X-0.412833333333 Y1.655 E1

G1 X-0.612 Y1.655 E1

G1 X-0.811166666667 Y1.655 E1

G1 X-1.01033333333 Y1.655 E1

G1 X-1.2095 Y1.655 E1

G1 X-1.40866666667 Y1.655 E1

G1 X-1.60783333333 Y1.655 E1

G1 X-1.807 Y1.655 E1

G1 X-2.00616666667 Y1.655 E1

G1 X-2.20533333333 Y1.655 E1

G1 X-2.4045 Y1.655 E1

G1 X-2.60366666667 Y1.655 E1

G1 X-2.80283333333 Y1.655 E1

G1 X-3.002 Y1.655 E0.12313 ; perimeter

G1 X-3.118 Y1.630 E0.00244 ; perimeter

G1 X-3.247 Y1.552 E0.00312 ; perimeter

G1 X-3.3155 Y1.479 E1

G1 X-3.384 Y1.406 E0.00412 ; perimeter

G1 X-3.4495 Y1.299 E1

G1 X-3.515 Y1.192 E0.00517 ; perimeter

G1 X-3.5705 Y1.057 E1

G1 X-3.626 Y0.922 E0.00602 ; perimeter

G1 X-3.66533333333 Y0.756666666667 E1

G1 X-3.70466666667 Y0.591333333333 E1

G1 X-3.744 Y0.426 E0.01049 ; perimeter

G1 X-3.75766666667 Y0.243 E1

G1 X-3.77133333333 Y0.06 E1

G1 X-3.785 Y-0.123 E0.01135 ; perimeter

G1 X-3.776 Y-0.3055 E1

G1 X-3.767 Y-0.488 E0.00752 ; perimeter

```
G1 X-3.7405 Y-0.6615 E1
G1 X-3.714 Y-0.835 E0.00725 ; perimeter
G1 X-3.66733333333 Y-0.990333333333 E1
G1 X-3.62066666667 Y-1.14566666667 E1
G1 X-3.574 Y-1.301 E0.01002 ; perimeter
G1 X-3.4775 Y-1.475 E1
G1 X-3.381 Y-1.649 E0.00820 ; perimeter
G1 X-3.284 Y-1.743 E1
G1 X-3.187 Y-1.837 E0.00556 ; perimeter
M761
G1 X-2.9045 Y-1.765 E1
G1 X-2.622 Y-1.693 F600.000 ; move to first infill point
M760
G4 P0
G1 X-2.816 Y-1.647 E0.00451 F240.000 ; infill
G1 X-2.917 Y-1.606 E1
G1 X-3.018 Y-1.565 E0.00490 ; infill
G1 X-3.120 Y-1.459 E0.00331 ; infill
G1 X-3.1875 Y-1.3535 E1
G1 X-3.255 Y-1.248 E0.00564 ; infill
G1 X-3.31433333333 Y-1.08933333333 E1
G1 X-3.37366666667 Y-0.930666666667 E1
G1 X-3.433 Y-0.772 E0.01144 ; infill
G1 X-3.461 Y-0.616 E1
G1 X-3.489 Y-0.460 E0.00714 ; infill
G1 X-3.498 Y-0.293 E1
G1 X-3.507 Y-0.126 E0.00751 ; infill
G1 X-3.499 Y0.026 E1
G1 X-3.491 Y0.178 E0.00686 ; infill
G1 X-3.4685 Y0.3235 E1
G1 X-3.446 Y0.469 E0.00662 ; infill
G1 X-3.404 Y0.618 E1
G1 X-3.362 Y0.767 E0.00696 ; infill
```

```
G1 X-3.302 Y0.903 E1
G1 X-3.242 Y1.039 E0.00669 ; infill
G1 X-3.185 Y1.1245 E1
G1 X-3.128 Y1.210 E0.00464 ; infill
G1 X-3.000 Y1.335 E0.00401 ; infill
G1 X-2.817 Y1.407 E0.00444 ; infill
G1 X-2.719 Y1.43 E1
G1 X-2.621 Y1.453 E0.00451 ; infill
M761
G1 E-2.00000 F2400.00000 ; retract
G1 X-2.14463636364 Y1.45290909091 E1
G1 X-1.66827272727 Y1.45281818182 E1
G1 X-1.19190909091 Y1.45272727273 E1
G1 X-0.715545454545 Y1.45263636364 E1
G1 X-0.239181818182 Y1.45254545455 E1
G1 X0.237181818182 Y1.45245454545 E1
G1 X0.713545454545 Y1.45236363636 E1
G1 X1.18990909091 Y1.45227272727 E1
G1 X1.66627272727 Y1.45218181818 E1
G1 X2.14263636364 Y1.45209090909 E1
G1 X2.619 Y1.452 F600.000 ; move to first infill point
M760
G4 P0
G1 X-0.874333333333 Y1.45266666667 E1
G1 X0.872333333333 Y1.45233333333 E1
G1 E2.00000 F2400.00000 ; unretract
G1 X2.746 Y1.40933333333 E1
G1 X2.873 Y1.36666666667 E1
G1 X3.000 Y1.324 E0.00904 F240.000 ; infill
G1 X3.108 Y1.210 E0.00352 ; infill
G1 X3.1675 Y1.121 E1
G1 X3.227 Y1.032 E0.00483 ; infill
G1 X3.2735 Y0.9265 E1
```

```
G1 X3.320 Y0.821 E0.00518 ; infill
G1 X3.3655 Y0.6805 E1
G1 X3.411 Y0.540 E0.00666 ; infill
G1 X3.442 Y0.367 E1
G1 X3.473 Y0.194 E0.00791 ; infill
G1 X3.4795 Y0.0345 E1
G1 X3.486 Y-0.125 E0.00716 ; infill
G1 X3.479 Y-0.2725 E1
G1 X3.472 Y-0.420 E0.00666 ; infill
G1 X3.441 Y-0.603 E1
G1 X3.410 Y-0.786 E0.00835 ; infill
G1 X3.365 Y-0.9235 E1
G1 X3.320 Y-1.061 E0.00651 ; infill
G1 X3.2735 Y-1.1665 E1
G1 X3.227 Y-1.272 E0.00518 ; infill
G1 X3.1675 Y-1.361 E1
G1 X3.108 Y-1.450 E0.00483 ; infill
G1 X2.984 Y-1.572 E0.00391 ; infill
G1 X2.89 Y-1.609 E1
G1 X2.796 Y-1.646 E0.00454 ; infill
G1 X2.603 Y-1.692 E0.00446 ; infill
M761
G1 E-2.00000 F2400.00000 ; retract
G0 Z30
M84 ;disable motors
;END
```

**Model 2 (interlocking rings)**

```
; external perimeters extrusion width = 0.45mm
; perimeters extrusion width = 0.48mm
; infill extrusion width = 0.48mm
; solid infill extrusion width = 0.48mm
; top infill extrusion width = 0.48mm
; support material extrusion width = 0.45mm
G21 ; set units to millimeters
G90 ; use absolute coordinates
M83 ; use relative distances for extrusion
G1 Z0.400 F4800.000 ; move to next layer (0)
G1 X0.067 Y0.292 F4800.000 ; move to first perimeter point
M760
G4 P0
G1 X0.133 Y0.512 E0.00408 F360.000 ; perimeter
G1 X-0.014 Y0.581 E0.00289 ; perimeter
G1 X-0.037 Y0.547 E0.00073 ; perimeter
G1 X-0.122 Y0.512 E0.00163 ; perimeter
G1 X-0.080 Y0.299 E0.00385 ; perimeter
G1 X-0.000 Y0.295 E0.00143 ; perimeter
M761
G1 X0.067 Y-0.292 F4800.000 ; move to first perimeter point
M760
G4 P0
G1 X-0.080 Y-0.299 E0.00263 F360.000 ; perimeter
G1 X-0.122 Y-0.512 E0.00385 ; perimeter
G1 X-0.037 Y-0.547 E0.00163 ; perimeter
G1 X-0.014 Y-0.581 E0.00072 ; perimeter
G1 X0.133 Y-0.512 E0.00289 ; perimeter
G1 X0.087 Y-0.357 E0.00288 ; perimeter
M761
G1 E-2.00000 F2400.00000 ; retract
G1 X-3.445 Y-0.172 F4800.000 ; move to first perimeter point
```

```
M760
G4 P0
G1 X-1.679 Y-0.2645 E1
G1 E2.00000 F2400.00000 ; unretract
G1 X-3.445 Y0.0 E1
G1 X-3.445 Y0.172 E0.00611 F360.000 ; perimeter
G1 X-3.4115 Y0.3405 E1
G1 X-3.378 Y0.509 E0.00611 ; perimeter
G1 X-3.313 Y0.668 E1
G1 X-3.248 Y0.827 E0.00611 ; perimeter
G1 X-3.1535 Y0.97 E1
G1 X-3.059 Y1.113 E0.00609 ; perimeter
G1 X-2.937 Y1.236 E1
G1 X-2.815 Y1.359 E0.00615 ; perimeter
G1 X-2.6675 Y1.4535 E1
G1 X-2.520 Y1.548 E0.00623 ; perimeter
G1 X-2.3625 Y1.612 E1
G1 X-2.205 Y1.676 E0.00605 ; perimeter
G1 X-2.0385 Y1.709 E1
G1 X-1.872 Y1.742 E0.00604 ; perimeter
G1 X-1.7025 Y1.74 E1
G1 X-1.533 Y1.738 E0.00603 ; perimeter
G1 X-1.249 Y1.676 E0.00517 ; perimeter
G1 X-1.064 Y1.612 E0.00347 ; perimeter
G1 X-0.921 Y1.547 E0.00279 ; perimeter
G1 X-0.7755 Y1.449 E1
G1 X-0.630 Y1.351 E0.00625 ; perimeter
G1 X-0.5145 Y1.2345 E1
G1 X-0.399 Y1.118 E0.00584 ; perimeter
G1 X-0.296 Y0.965 E0.00327 ; perimeter
G1 X-0.078 Y0.957 E0.00388 ; perimeter
G1 X0.063 Y0.923 E0.00257 ; perimeter
G1 X0.292 Y0.964 E0.00414 ; perimeter
```

```
G1 X0.469 Y1.193 E0.00515 ; perimeter
G1 X0.608 Y1.3135 E1
G1 X0.747 Y1.434 E0.00654 ; perimeter
G1 X0.918 Y1.541 E0.00357 ; perimeter
G1 X1.0735 Y1.606 E1
G1 X1.229 Y1.671 E0.00599 ; perimeter
G1 X1.413 Y1.715 E0.00337 ; perimeter
G1 X1.702 Y1.745 E0.00516 ; perimeter
G1 X1.894 Y1.737 E0.00342 ; perimeter
G1 X2.0595 Y1.704 E1
G1 X2.225 Y1.671 E0.00600 ; perimeter
G1 X2.410 Y1.601 E0.00351 ; perimeter
G1 X2.666 Y1.462 E0.00518 ; perimeter
G1 X2.817 Y1.352 E0.00333 ; perimeter
G1 X2.9365 Y1.2315 E1
G1 X3.056 Y1.111 E0.00603 ; perimeter
G1 X3.172 Y0.947 E0.00357 ; perimeter
G1 X3.2455 Y0.812 E1
G1 X3.319 Y0.677 E0.00546 ; perimeter
G1 X3.3695 Y0.507 E1
G1 X3.420 Y0.337 E0.00630 ; perimeter
G1 X3.4365 Y0.1715 E1
G1 X3.453 Y0.006 E0.00593 ; perimeter
G1 X3.436 Y-0.1685 E1
G1 X3.419 Y-0.343 E0.00623 ; perimeter
G1 X3.369 Y-0.51 E1
G1 X3.319 Y-0.677 E0.00620 ; perimeter
G1 X3.2325 Y-0.832 E1
G1 X3.146 Y-0.987 E0.00630 ; perimeter
G1 X3.038 Y-1.1175 E1
G1 X2.930 Y-1.248 E0.00603 ; perimeter
G1 X2.706 Y-1.435 E0.00518 ; perimeter
G1 X2.5485 Y-1.525 E1
```

```
G1 X2.391 Y-1.615 E0.00645 ; perimeter
G1 X2.2245 Y-1.666 E1
G1 X2.058 Y-1.717 E0.00619 ; perimeter
G1 X1.8915 Y-1.731 E1
G1 X1.725 Y-1.745 E0.00594 ; perimeter
G1 X1.5615 Y-1.731 E1
G1 X1.398 Y-1.717 E0.00584 ; perimeter
G1 X1.22 Y-1.661 E1
G1 X1.042 Y-1.605 E0.00663 ; perimeter
G1 X0.788 Y-1.462 E0.00518 ; perimeter
G1 X0.632 Y-1.348 E0.00344 ; perimeter
G1 X0.488 Y-1.217 E0.00347 ; perimeter
G1 X0.39 Y-1.0905 E1
G1 X0.292 Y-0.964 E0.00569 ; perimeter
G1 X0.063 Y-0.923 E0.00414 ; perimeter
G1 X-0.078 Y-0.957 E0.00257 ; perimeter
G1 X-0.296 Y-0.965 E0.00388 ; perimeter
G1 X-0.395 Y-1.113 E0.00317 ; perimeter
G1 X-0.5145 Y-1.234 E1
G1 X-0.634 Y-1.355 E0.00605 ; perimeter
G1 X-0.775 Y-1.4495 E1
G1 X-0.916 Y-1.544 E0.00603 ; perimeter
G1 X-1.0715 Y-1.6095 E1
G1 X-1.227 Y-1.675 E0.00601 ; perimeter
G1 X-1.393 Y-1.708 E1
G1 X-1.559 Y-1.741 E0.00601 ; perimeter
G1 X-1.847 Y-1.743 E0.00511 ; perimeter
G1 X-2.037 Y-1.709 E1
G1 X-2.227 Y-1.675 E0.00686 ; perimeter
G1 X-2.3955 Y-1.6015 E1
G1 X-2.564 Y-1.528 E0.00653 ; perimeter
G1 X-2.692 Y-1.4415 E1
G1 X-2.820 Y-1.355 E0.00549 ; perimeter
```

```
G1 X-2.9395 Y-1.234 E1
G1 X-3.059 Y-1.113 E0.00605 ; perimeter
G1 X-3.1595 Y-0.9565 E1
G1 X-3.260 Y-0.800 E0.00661 ; perimeter
G1 X-3.319 Y-0.6545 E1
G1 X-3.378 Y-0.509 E0.00559 ; perimeter
G1 X-3.4115 Y-0.3405 E1
G1 X-3.445 Y-0.172 E0.00611 ; perimeter
M761
G1 Z0.800 F4800.000 ; move to next layer (1)
G1 E-2.00000 F2400.00000 ; retract
G1 X0.288 Y0.321 F4800.000 ; move to first perimeter point
M760
G4 P0
G1 X-1.5785 Y0.0745 E1
G1 E2.00000 F2400.00000 ; unretract
G1 X0.128666666667 Y0.481333333333 E1
G1 X-0.0306666666667 Y0.641666666667 E1
G1 X-0.190 Y0.802 E0.01526 F360.000 ; perimeter
G1 X-0.2945 Y0.96 E1
G1 X-0.399 Y1.118 E0.00851 ; perimeter
G1 X-0.512 Y1.238 E0.00371 ; perimeter
G1 X-0.630 Y1.351 E0.00368 ; perimeter
G1 X-0.76 Y1.4395 E1
G1 X-0.890 Y1.528 E0.00709 ; perimeter
G1 X-1.064 Y1.612 E0.00434 ; perimeter
G1 X-1.231 Y1.663 E1
G1 X-1.398 Y1.714 E0.00786 ; perimeter
G1 X-1.553 Y1.741 E0.00353 ; perimeter
G1 X-1.727 Y1.741 E1
G1 X-1.901 Y1.741 E0.00783 ; perimeter
G1 X-2.077 Y1.7015 E1
G1 X-2.253 Y1.662 E0.00811 ; perimeter
```

G1 X-2.520 Y1.548 E0.00655 ; perimeter

G1 X-2.6675 Y1.4535 E1

G1 X-2.815 Y1.359 E0.00789 ; perimeter

G1 X-2.937 Y1.236 E1

G1 X-3.059 Y1.113 E0.00778 ; perimeter

G1 X-3.1535 Y0.97 E1

G1 X-3.248 Y0.827 E0.00771 ; perimeter

G1 X-3.313 Y0.668 E1

G1 X-3.378 Y0.509 E0.00774 ; perimeter

G1 X-3.4115 Y0.3405 E1

G1 X-3.445 Y0.172 E0.00773 ; perimeter

G1 X-3.445 Y0.0125 E1

G1 X-3.445 Y-0.147 E0.00717 ; perimeter

G1 X-3.4075 Y-0.338 E1

G1 X-3.370 Y-0.529 E0.00877 ; perimeter

G1 X-3.309 Y-0.678 E1

G1 X-3.248 Y-0.827 E0.00725 ; perimeter

G1 X-3.1475 Y-0.976 E1

G1 X-3.047 Y-1.125 E0.00809 ; perimeter

G1 X-2.9335 Y-1.24 E1

G1 X-2.820 Y-1.355 E0.00727 ; perimeter

G1 X-2.679 Y-1.4495 E1

G1 X-2.538 Y-1.544 E0.00763 ; perimeter

G1 X-2.3825 Y-1.6095 E1

G1 X-2.227 Y-1.675 E0.00760 ; perimeter

G1 X-2.061 Y-1.708 E1

G1 X-1.895 Y-1.741 E0.00761 ; perimeter

G1 X-1.727 Y-1.741 E1

G1 X-1.559 Y-1.741 E0.00756 ; perimeter

G1 X-1.4035 Y-1.71 E1

G1 X-1.248 Y-1.679 E0.00714 ; perimeter

G1 X-1.069 Y-1.6035 E1

G1 X-0.890 Y-1.528 E0.00874 ; perimeter

```
G1 X-0.762 Y-1.4415 E1
G1 X-0.634 Y-1.355 E0.00695 ; perimeter
G1 X-0.5205 Y-1.24 E1
G1 X-0.407 Y-1.125 E0.00727 ; perimeter
G1 X-0.294 Y-0.957 E1
G1 X-0.181 Y-0.789 E0.00913 ; perimeter
G1 X-0.0243333333333 Y-0.634666666667 E1
G1 X0.132333333333 Y-0.480333333333 E1
G1 X0.289 Y-0.326 E0.01484 ; perimeter
M761
G1 X0.000 Y-0.197 F4800.000 ; move to first perimeter point
M760
G4 P0
G1 X0.0 Y0.0723333333333 E1
G1 X0.0 Y0.341666666667 E1
G1 X-0.000 Y0.611 E0.01816 F360.000 ; perimeter
G1 X0.190 Y0.802 E0.00607 ; perimeter
G1 X0.2925 Y0.9575 E1
G1 X0.395 Y1.113 E0.00838 ; perimeter
G1 X0.5145 Y1.234 E1
G1 X0.634 Y1.355 E0.00766 ; perimeter
G1 X0.775 Y1.4495 E1
G1 X0.916 Y1.544 E0.00763 ; perimeter
G1 X1.0715 Y1.6095 E1
G1 X1.227 Y1.675 E0.00760 ; perimeter
G1 X1.393 Y1.708 E1
G1 X1.559 Y1.741 E0.00761 ; perimeter
G1 X1.727 Y1.741 E1
G1 X1.895 Y1.741 E0.00756 ; perimeter
G1 X2.061 Y1.708 E1
G1 X2.227 Y1.675 E0.00761 ; perimeter
G1 X2.3825 Y1.6095 E1
G1 X2.538 Y1.544 E0.00760 ; perimeter
```

G1 X2.679 Y1.4495 E1

G1 X2.820 Y1.355 E0.00763 ; perimeter

G1 X2.9395 Y1.234 E1

G1 X3.059 Y1.113 E0.00765 ; perimeter

G1 X3.155 Y0.9675 E1

G1 X3.251 Y0.822 E0.00785 ; perimeter

G1 X3.3095 Y0.674 E1

G1 X3.368 Y0.526 E0.00715 ; perimeter

G1 X3.4065 Y0.349 E1

G1 X3.445 Y0.172 E0.00816 ; perimeter

G1 X3.4415 Y-0.011 E1

G1 X3.438 Y-0.194 E0.00823 ; perimeter

G1 X3.403 Y-0.361 E1

G1 X3.368 Y-0.528 E0.00769 ; perimeter

G1 X3.3095 Y-0.675 E1

G1 X3.251 Y-0.822 E0.00711 ; perimeter

G1 X3.155 Y-0.9675 E1

G1 X3.059 Y-1.113 E0.00785 ; perimeter

G1 X2.9395 Y-1.234 E1

G1 X2.820 Y-1.355 E0.00765 ; perimeter

G1 X2.6765 Y-1.451 E1

G1 X2.533 Y-1.547 E0.00777 ; perimeter

G1 X2.369 Y-1.6115 E1

G1 X2.205 Y-1.676 E0.00792 ; perimeter

G1 X2.0405 Y-1.708 E1

G1 X1.876 Y-1.740 E0.00755 ; perimeter

G1 X1.7155 Y-1.7385 E1

G1 X1.555 Y-1.737 E0.00721 ; perimeter

G1 X1.378 Y-1.6995 E1

G1 X1.201 Y-1.662 E0.00814 ; perimeter

G1 X1.061 Y-1.6045 E1

G1 X0.921 Y-1.547 E0.00681 ; perimeter

G1 X0.7755 Y-1.449 E1

```
G1 X0.630 Y-1.351 E0.00791 ; perimeter
G1 X0.5145 Y-1.2345 E1
G1 X0.399 Y-1.118 E0.00740 ; perimeter
G1 X0.2945 Y-0.96 E1
G1 X0.190 Y-0.802 E0.00851 ; perimeter
G1 X0.000 Y-0.611 E0.00606 ; perimeter
G1 X0.0 Y-0.404 E1
G1 X0.000 Y-0.197 E0.00931 ; perimeter
M761
G1 Z1.200 F4800.000 ; move to next layer (2)
G1 X0.297 Y0.364 F4800.000 ; move to first perimeter point
M760
G4 P0
G1 X0.093 Y0.535 E1
G1 X-0.111 Y0.706 E0.01199 F360.000 ; perimeter
G1 X-0.236666666667 Y0.876333333333 E1
G1 X-0.362333333333 Y1.04666666667 E1
G1 X-0.488 Y1.217 E0.01427 ; perimeter
G1 X-0.638 Y1.3395 E1
G1 X-0.788 Y1.462 E0.00873 ; perimeter
G1 X-0.9255 Y1.5385 E1
G1 X-1.063 Y1.615 E0.00707 ; perimeter
G1 X-1.2305 Y1.666 E1
G1 X-1.398 Y1.717 E0.00787 ; perimeter
G1 X-1.55 Y1.731 E1
G1 X-1.702 Y1.745 E0.00688 ; perimeter
G1 X-1.901 Y1.737 E0.00447 ; perimeter
G1 X-2.056 Y1.717 E0.00353 ; perimeter
G1 X-2.2335 Y1.6605 E1
G1 X-2.411 Y1.604 E0.00839 ; perimeter
G1 X-2.5595 Y1.521 E1
G1 X-2.708 Y1.438 E0.00765 ; perimeter
G1 X-2.872 Y1.286 E1
```

```
G1 X-3.036 Y1.134 E0.01005 ; perimeter
G1 X-3.1405 Y0.979 E1
G1 X-3.245 Y0.824 E0.00841 ; perimeter
G1 X-3.327 Y0.644 E0.00445 ; perimeter
G1 X-3.3735 Y0.48 E1
G1 X-3.420 Y0.316 E0.00767 ; perimeter
G1 X-3.434 Y0.146 E1
G1 X-3.448 Y-0.024 E0.00768 ; perimeter
G1 X-3.420 Y-0.316 E0.00660 ; perimeter
G1 X-3.3735 Y-0.48 E1
G1 X-3.327 Y-0.644 E0.00767 ; perimeter
G1 X-3.2495 Y-0.7955 E1
G1 X-3.172 Y-0.947 E0.00766 ; perimeter
G1 X-3.0675 Y-1.0805 E1
G1 X-2.963 Y-1.214 E0.00763 ; perimeter
G1 X-2.817 Y-1.352 E0.00451 ; perimeter
G1 X-2.6765 Y-1.4465 E1
G1 X-2.536 Y-1.541 E0.00761 ; perimeter
G1 X-2.3805 Y-1.606 E1
G1 X-2.225 Y-1.671 E0.00759 ; perimeter
G1 X-2.06 Y-1.7045 E1
G1 X-1.895 Y-1.738 E0.00760 ; perimeter
G1 X-1.7275 Y-1.7375 E1
G1 X-1.560 Y-1.737 E0.00754 ; perimeter
G1 X-1.366 Y-1.706 E0.00442 ; perimeter
G1 X-1.205 Y-1.6535 E1
G1 X-1.044 Y-1.601 E0.00761 ; perimeter
G1 X-0.788 Y-1.462 E0.00655 ; perimeter
G1 X-0.6395 Y-1.338 E1
G1 X-0.491 Y-1.214 E0.00871 ; perimeter
G1 X-0.364333333333 Y-1.04466666667 E1
G1 X-0.237666666667 Y-0.875333333333 E1
G1 X-0.111 Y-0.706 E0.01427 ; perimeter
```

```
G1 X0.093 Y-0.535 E1
G1 X0.297 Y-0.364 E0.01199 ; perimeter
M761
G1 X-0.003 Y-0.197 F4800.000 ; move to first perimeter point
M760
G4 P0
G1 X-0.00233333333333 Y0.072 E1
G1 X-0.00166666666667 Y0.341 E1
G1 X-0.001 Y0.610 E0.01817 F360.000 ; perimeter
G1 X0.190 Y0.802 E0.00609 ; perimeter
G1 X0.2925 Y0.9575 E1
G1 X0.395 Y1.113 E0.00838 ; perimeter
G1 X0.5145 Y1.234 E1
G1 X0.634 Y1.355 E0.00766 ; perimeter
G1 X0.775 Y1.4495 E1
G1 X0.916 Y1.544 E0.00763 ; perimeter
G1 X1.0715 Y1.6095 E1
G1 X1.227 Y1.675 E0.00760 ; perimeter
G1 X1.393 Y1.708 E1
G1 X1.559 Y1.741 E0.00761 ; perimeter
G1 X1.727 Y1.741 E1
G1 X1.895 Y1.741 E0.00756 ; perimeter
G1 X2.061 Y1.708 E1
G1 X2.227 Y1.675 E0.00761 ; perimeter
G1 X2.3825 Y1.6095 E1
G1 X2.538 Y1.544 E0.00760 ; perimeter
G1 X2.679 Y1.4495 E1
G1 X2.820 Y1.355 E0.00763 ; perimeter
G1 X2.9395 Y1.234 E1
G1 X3.059 Y1.113 E0.00765 ; perimeter
G1 X3.155 Y0.9675 E1
G1 X3.251 Y0.822 E0.00785 ; perimeter
G1 X3.316 Y0.6525 E1
```

G1 X3.381 Y0.483 E0.00817 ; perimeter

G1 X3.413 Y0.3275 E1

G1 X3.445 Y0.172 E0.00714 ; perimeter

G1 X3.4415 Y-0.011 E1

G1 X3.438 Y-0.194 E0.00823 ; perimeter

G1 X3.403 Y-0.361 E1

G1 X3.368 Y-0.528 E0.00769 ; perimeter

G1 X3.3095 Y-0.675 E1

G1 X3.251 Y-0.822 E0.00711 ; perimeter

G1 X3.155 Y-0.9675 E1

G1 X3.059 Y-1.113 E0.00785 ; perimeter

G1 X2.9395 Y-1.234 E1

G1 X2.820 Y-1.355 E0.00765 ; perimeter

G1 X2.6765 Y-1.451 E1

G1 X2.533 Y-1.547 E0.00777 ; perimeter

G1 X2.369 Y-1.6115 E1

G1 X2.205 Y-1.676 E0.00792 ; perimeter

G1 X1.920 Y-1.734 E0.00654 ; perimeter

G1 X1.7635 Y-1.7365 E1

G1 X1.607 Y-1.739 E0.00705 ; perimeter

G1 X1.4215 Y-1.706 E1

G1 X1.236 Y-1.673 E0.00847 ; perimeter

G1 X1.0785 Y-1.61 E1

G1 X0.921 Y-1.547 E0.00763 ; perimeter

G1 X0.7755 Y-1.449 E1

G1 X0.630 Y-1.351 E0.00791 ; perimeter

G1 X0.5145 Y-1.2345 E1

G1 X0.399 Y-1.118 E0.00740 ; perimeter

G1 X0.2945 Y-0.96 E1

G1 X0.190 Y-0.802 E0.00851 ; perimeter

G1 X-0.001 Y-0.610 E0.00609 ; perimeter

G1 X-0.002 Y-0.4035 E1

G1 X-0.003 Y-0.197 E0.00929 ; perimeter

```
M761
G1 E-2.00000 F2400.00000 ; retract
G0 Z30
M84 ;disable motors
;END
```
